# FASTIMAGES: Validating replay detection methods in human neuroimaging using a combined MEG and fMRI dataset

**DOI:** 10.64898/2026.05.26.727586

**Authors:** Simon Kern, Lennart Wittkuhn, Elena Buss, Nicolas Schuck, Gordon B. Feld

## Abstract

Studies in rodents and humans using invasive electrophysiology have established that neural replay is a ubiquitous phenomenon in the brain that is associated with a wide range of cognitive functions, including memory, planning and decision making. Yet, invasively recording in humans remains difficult, and hence knowledge about replay in humans remains scarce. Hence, to comprehensively understand replay in humans, we need reliable approaches that can detect it non-invasively. Several main non-invasive approaches have been proposed, but we lack a full comparative validation against known ground truth signals. In this study, we present FASTIMAGES, a benchmark dataset from seventy participants with parallel fMRI (n = 40, previously published) and MEG (n=30) recordings containing known neural sequences evoked by fast visual stimulation as well as functional localizer trials. The neural sequences were elicited by five different visual stimuli shown in sequences at speeds of 132, 164, 228 and 612 milliseconds onset-to-onset intervals. Using this dataset, we investigate two existing statistical methods for sequence detection, namely Temporally Delayed Linear Modelling (TDLM, developed for MEG by Liu et al., 2021) and Slope Order Dynamic Analysis (SODA, developed for fMRI by Wittkuhn & Schuck, 2021). We examine the underlying assumptions of each method, analyse their resulting strengths and weaknesses in application to MEG and fMRI. We demonstrate that both approaches excel in their native modality (TDLM for MEG and SODA for fMRI), with comparable effect sizes given idealized conditions in this benchmark. Cross-modality transfer remains challenging. Finally, the FASTIMAGES dataset provides data with known and clearly expressed sequences and can be used to benchmark and validate future sequence detection methods under idealized conditions.

## Introduction

Neural replay of past experiences has emerged as a widespread phenomenon hypothesized to be involved in many different cognitive processes, most prominently decision making, consolidation and planning (Born & Wilhelm, 2012; Carr et al., 2011; Feld & Born, 2017; Foster, 2017). “Replay” is generally defined as roughly temporally preserved sequences of neuronal activation patterns that occur without external stimulation but recapitulate previously encountered situations (Genzel et al., 2020). Replay more specifically denotes a re-appearance of a neuronal pattern in the absence of the original percept, following the same temporal pattern as during learning of the information (Z. S. Chen & Wilson, 2023). Typically, these replay events are time-compressed (Nádasdy et al., 1999) and may unfold in the experienced (forward) or inverted (reverse) order (Ambrose et al., 2016; Diba & Buzsáki, 2007; Foster & Wilson, 2006). Although first characterized in rodent hippocampal place cells (Nádasdy et al., 1999; Skaggs & McNaughton, 1996), sequential replay has since been documented across many mammalian species (Hoffman & McNaughton, 2002; Lee & Wilson, 2002; Pavlides & Winson, 1989) and, with intracranial recordings, in humans (Eichenlaub et al., 2020; Rubin et al., 2022; Vaz et al., 2020). Replay occurs mainly during offline states (Chang et al., 2025), but also during task pauses (Ólafsdóttir et al., 2016) and active performance (Ólafsdóttir et al., 2018). It stabilises memory traces through cortical plasticity (Diekelmann & Born, 2010), integrates experience with existing schemas to support abstraction (Lewis & Durrant, 2011; Tse et al., 2007), and enables planning by simulating future trajectories (Foster & Wilson, 2006; Pfeiffer & Foster, 2013) thus bridging past experience with future behaviour (Foster, 2017; Wittkuhn et al., 2021).

Two statistical methods have been proposed to detect sequential replay signatures non-invasively in human participants in recent years. Both methods rely on brain decoders to assess similarity of the current brain state to brain activity during a localizer task. “Temporally Delayed Linear Modelling” (TDLM, Liu et al., 2021) was initially developed in the domain of MEG and quantifies the temporal succession of individual states within neural data by testing whether pairwise temporal state successions follow an expected order more often than expected by chance (Figure 1 upper panels). Although validation of TDLM against empirical ground truth data has not been provided so far, it has been used in more than 20 different studies to date with an accumulated 1400+ citations (crossref), showing promise to detect replay, e.g., during planning and decision making (Eldar et al., 2020; McFadyen et al., 2023; Schwartenbeck et al., 2023; Wimmer et al., 2023; Wise et al., 2021), memory retrieval (Huang & Luo, 2023; Kern et al., 2024; Wimmer et al., 2020) or in the context of reward learning (Liu, Mattar, et al., 2021). A contribution of this paper is therefore to provide an empirical ground truth validation of TDLM. A second method that was introduced in the context of functional magnetic resonance imaging (fMRI) is “Slope Order Dynamic Analysis” (SODA, Schuck & Niv, 2019; Wittkuhn & Schuck, 2021). SODA quantifies whether reactivations are ordered by a specific gradient *within* each time point and then tracks how that gradient evolves over time to infer stimulus sequence, making use of the largely overlapping haemodynamic responses (in the time course of seconds) that follow a specific onset and offset dynamic. It was built on a predecessor method (Schuck & Niv, 2019) and has been used to demonstrate replay in the context of value learning (Renz et al., 2026), in the visual cortex in humans (Wittkuhn et al., 2025) and during resting state (Huang et al., 2024) (Figure 1 bottom panels).

**Figure 1.**
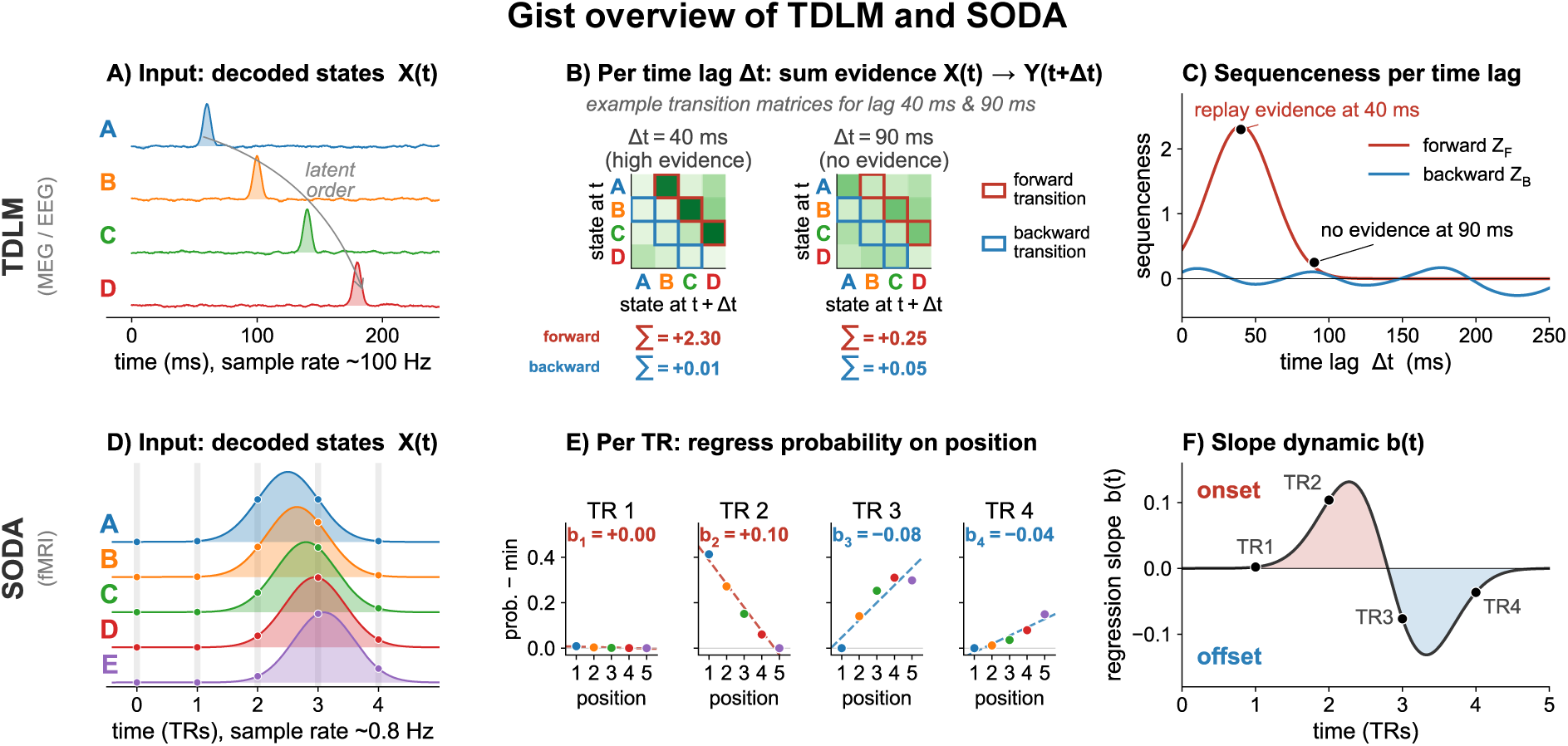
Visualization of TDLM and SODA, methods for neuronal sequence detection within their native measurement modality. **A)** TDLM works similarly to a Granger causality, assessing how much a specific reactivation predicts another reactivation at a later time step. Probabilities for reactivation of individual items (A, B, C, D) are tracked and expected to be temporally separate and brief, with a constant time lag during replay. **B)** For each time lag, a first-level GLM is used to construct an empirical weighted transition matrix: How much does a reactivation of item A at time t predict a reactivation of B at a later time lag? The evidence for the expected transitions as defined by the sequence is summed (red for forward, blue for backward transitions), yielding one “sequenceness” score for each time lag. **C)** The resulting sequenceness score tells how much evidence there is for temporal sequentiality at a specific time lag for the input sequence. **D)** SODA works by tracking reactivations that were in a specific rapid temporal order, but are largely overlapping in their measured modality (sampling at each TR, curves are fitted for visualization). **E)** For each TR, the state probabilities are regressed onto the target sequence, resulting in a slope per TR. Slopes are traditionally inverted. **F)** The change in slopes is analysed over time. If a prominent onset and offset phase is visible, there is evidence for replay. Frequency analysis of the slope can be used to estimate replay speed. NB: While previously called “forward” and “backward” period (Wittkuhn & Schuck, 2021), to avoid confusion with forward and backward replay, we will refer to them as “onset” and “offset” period in this paper.

A fundamental challenge in validating these statistical methods is the lack of ground truth, i.e., it is unknown how much – if any – replay occurs during a time window of interest. Such validation is essential because replay signals are typically small in magnitude while the analysis pipelines used to detect them are complex and offer many researcher degrees of freedom. This forking-path problem leads to inflated false-positive rates and undermines replicability. Simulations of MEG resting state data with “implanted” replay sequences (Kern et al., 2026; cf. Wittkuhn & Schuck, 2021) have shown that relatively high amounts of replay must be introduced when using TDLM (Kern et al., 2026). Moreover, our previous work has also shown that a common argument for TDLM’s validity - that replay strength is linked to behavioural performance (Wimmer et al., 2020) could reflect spurious effects that are not always reliable (Kern et al., 2026). These partially synthetic simulations do not necessarily capture the complex nature of neurophysiological recordings, and recreating naturalistic conditions remains challenging, so they constrain rather than settle the question of TDLM’s sensitivity in real data.

An alternative to simulations is to use a proxy task that evokes replay-like brain activity. Wittkuhn and Schuck (2021) developed such a task, in which fast visual sequences of five images were shown to participants. The visually evoked activity of each image creates a ground-truth dataset of known sequences which was used to validate the SODA algorithm. Here, we extend this work in two ways: we provide a novel MEG dataset acquired with the same task and perform new analyses of the 3T fMRI data from Wittkuhn and Schuck (2021). In what follows, we first present a harmonized MEG-fMRI dataset that can be used to validate replay detection pipelines, and then comparatively evaluate TDLM and SODA on this benchmark using bootstrap power analyses and effect size estimates. Additionally, we discuss problems in the cross-over application of SODA to MEG and TDLM to fMRI as well as potential solutions.

## Results Section 1 - Dataset

### FASTIMAGES - A sequence detection benchmark dataset

To achieve our aim of providing a comparative validation test, we acquired MEG data of 30 participants that contain fast activation sequences with known ground-truth, paralleling an existing fMRI dataset (Wittkuhn & Schuck, 2021). In addition to providing initial validation analyses, we release the harmonized joint data set, FASTIMAGES, as a resource for the community to facilitate further methodological developments.

In total, FASTIMAGES contains data from seventy healthy participants (thirty for the MEG, forty for the fMRI) observing visual images from five categories (house, cat, chair, face or shoe) either in a localizer or sequence condition (see Figure 2 for an overview). The localizer task involved the presentation of individual grayscale images for 500 ms and required participants to report when a shown image was upside down or not (as an attention check). To prevent any sequential bias, the order of localizer trials was pseudo-randomized, ensuring that all pairwise transitions were equally frequent, but excluding category repetitions. In the sequence condition, participants observed a sequence of five images (one from each category) for a short period in quick succession (32, 64, 128, 512 or 2048 milliseconds between offset of the previous image and onset of the next, ISI). Because each image was shown for 100 ms, the effective distance between two successive image onsets is 100 ms plus the ISI of 32 to 512 milliseconds. To ensure participants processed these sequences attentively, each trial started with a written category cue (‘shoe’), and ended with a serial order decision where participants needed to indicate at which position the cue image appeared (e.g., “3 or 5?”) The task and fMRI-data were previously reported in Wittkuhn & Schuck (2021), which we augment with a version of the task adapted to the MEG. See Methods section for details and differences between the fMRI and MEG task that were introduced for optimal measurement in each modality.

**Figure 2.**
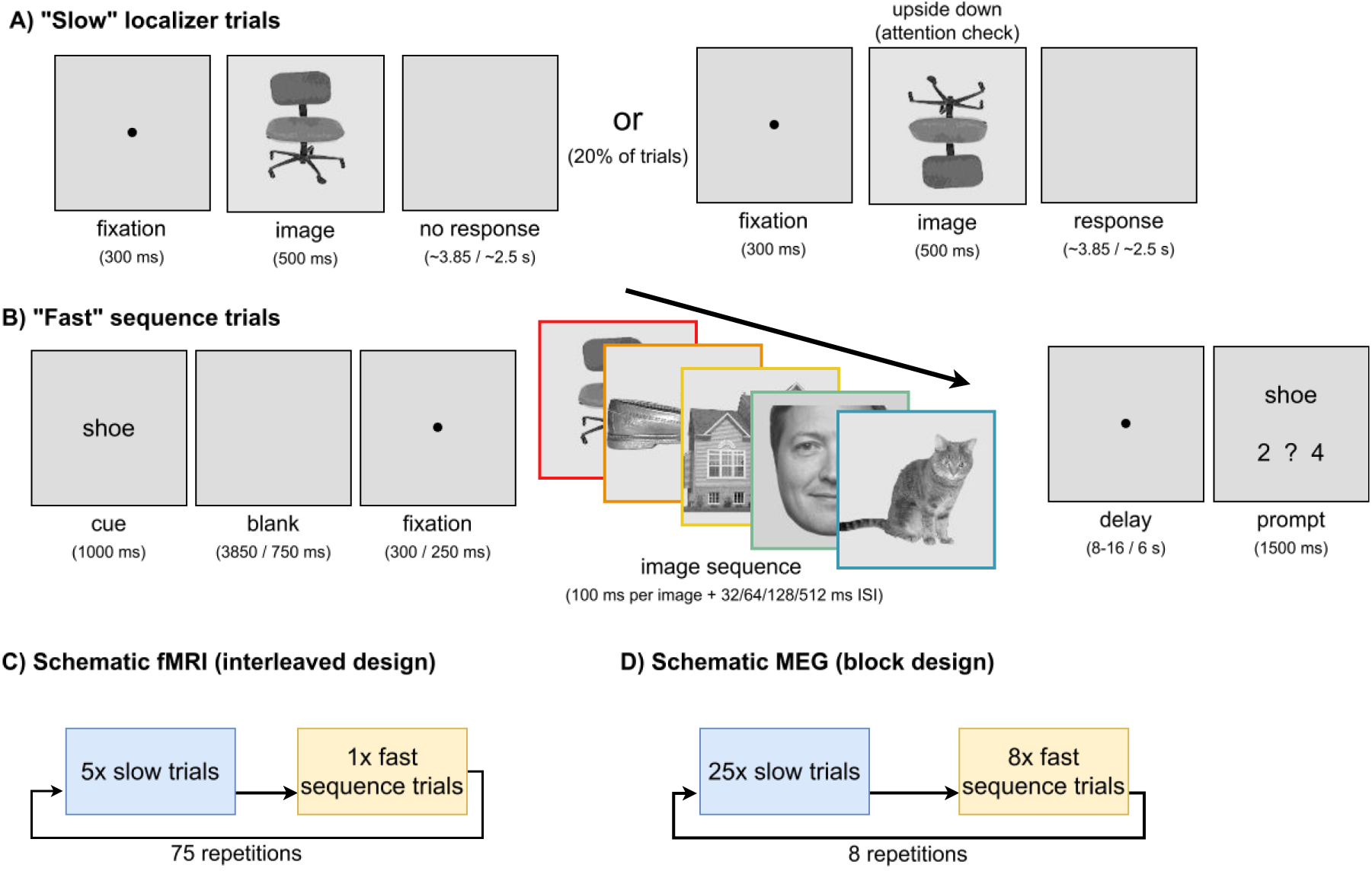
Schematic of the two types of paradigms recorded in the experiment and the differences between fMRI and MEG task implementation. **A)** “Slow” trials localizer: One of five grayscale images was shown for 500 ms, after which an inter-trial-interval (ITI) was blank. As an attention check, in ∼20% of cases the image was upside down, in which case the participant had to press a button as fast as possible. These trials were excluded from further analysis. **B)** “Fast” sequence trials: First a text cue describing one of the five images was shown and participants were asked to remember the position of the corresponding image in the upcoming sequence. Then, after an interval, a rapid sequence of five images was shown, with a different inter-stimulus-interval (ISI) per sequence. Each image within the sequence was shown for 100 ms, after which an ISI of 32, 64, 128, 512 or 2048 (fMRI only) was blank until the next image. Then, after a delay period, a binary choice had to be made between two positions corresponding to the position of the cued image. **C)** Implementation schematic of the fMRI task: The task always showed five localizer trials and one sequence trial interleaved, together with another type of trial (repetition trial, see Wittkuhn & Schuck, 2021, which is not part of this report). D) Implementation schematic of the MEG task: The task showed 25 localizer trials followed by eight sequence trials. Timings are shown in brackets, if two timings are indicated they are showing fMRI/MEG value pairs. Note: The image of the face in B) was replaced with a computer generated facsimile in accordance with Bioarxiv data protection rules.

### Localizer decoding

To demonstrate the category decodability of the localizer data, we assessed cross-validated performance of logistic regressions trained on each time point during the localizer trials separately. As shown in Figure 3A, decoding accuracy in both cases clearly surpassed chance level, with average peak decoding accuracy of ∼55% in the MEG (Figure 3A upper, chance level 20%, p<0.0001, t=14.97) and ∼70% in the fMRI (Figure 3A lower, chance level 20%, p<0.0001, t=21.35, see Supplement Figure 3 for category specific decoding probabilities of the five categories in MEG and fMRI, and subject level MEG decoding, respectively). Peak fMRI decoding surpassed MEG levels (p<0.0001, t=4.26); MEG decoding reached its peak around 150 milliseconds after stimulus onset with a rapid ramp-up and slow decay, fMRI decoding reflected the slower hemodynamic processes and peaked around four seconds. fMRI decoding accuracy was below chance at stimulus onset, likely reflecting activity of previous trials that were biased toward a different image than the current trial (images could not repeat). Final decoders for all subsequently shown analyses were trained on the time point of the peak average decoding accuracy, rounded to the nearest TR for the fMRI (see Methods section for details).

**Figure 3.**
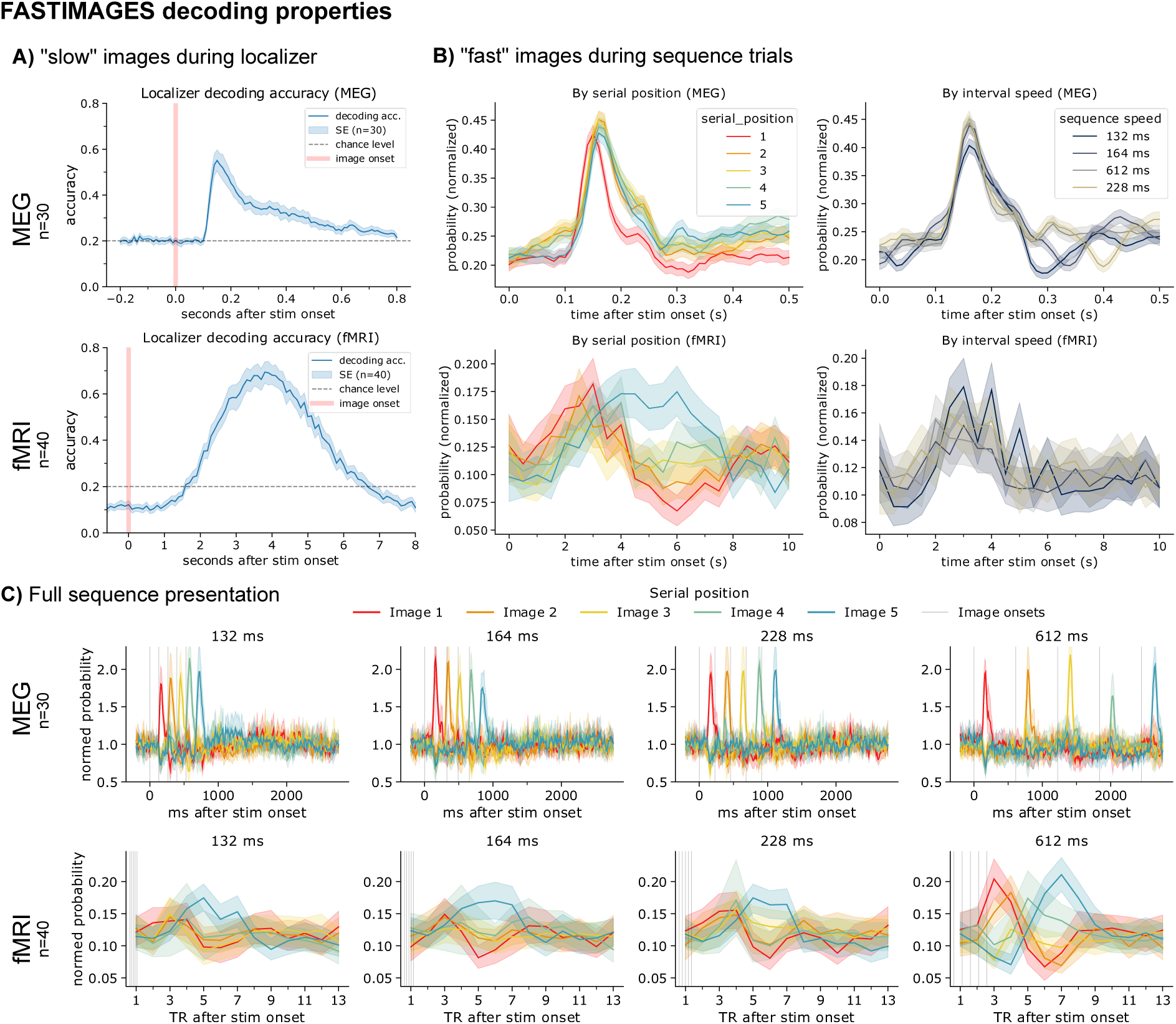
FASTIMAGES decoding properties: Decoding of the “slow” (localizer) and “fast” sequence trials for the MEG and fMRI conditions. **A)** Decoding accuracy during the localizer images. Cross-validation was used to test and train how well images could be predicted by the brain activity at that individual time point using a logistic regression. Final decoders were trained on the timepoint of peak decoding accuracy across all participants, rounded to the nearest TR for the fMRI trials. **B)** Mean decoded probability estimates (normalized) from a decoder trained on the localizer task (see A) during the sequence presentation, aligned on each individual image onset during the sequence for the MEG (upper) and fMRI (lower). Left: Sorted by serial position. Right: Sorted by presentation speed. **C)** Mean decoded probability estimates for the sequence task in which the five images were shown in fast succession, by a decoder trained on the localizer (see A). The stimuli were randomly distributed to sequence positions across participants. In the fMRI an additional slower condition was recorded, which used an onset-to-onset of 2148 ms, which we omitted. Shown are averages per participant with grey lines denoting the individual image onsets and bands denoting the standard error. fMRI timings have been rounded to the nearest 0.5 seconds. See https://github.com/CIMH-Clinical-Psychology/FASTIMAGES-benchmark/ for instructions on how to download the dataset.

### Sequence decoding

Individual image processing during the sequence presentation can be inspected by superimposing time series relative to each image onset (Figure 3B). In the MEG (Figure 3B upper), and fMRI (Figure 3B lower), images are well decodable above chance, both when split up by serial position and sequence speed. Decoding traces across sequence speeds appear broadly similar in both modalities, and decoding traces across serial positions also appear broadly similar.

Next, we used the FASTIMAGES dataset to benchmark the sequence detection abilities of TDLM and SODA. We first decoded the image probabilities from the time series data during the presentation of the sequences (Figure 3C), sorted by their presentation order. In MEG data (Figure 3C upper), sequences are distinctly visible in average plots even in the fastest condition (132 milliseconds onset-to-onset interval), with probabilities rapidly declining as soon as the next image of the sequence appears (Supplement Figure 6). In fMRI data (Figure 3C lower), the full sequence is visible by eye in the 612-millisecond condition, while only the order of the first and last stimuli is easily recognizable in the fastest condition.

We provide the probability time series for both the MEG and fMRI at GitHub at: https://github.com/CIMH-Clinical-Psychology/FASTIMAGES-benchmark/

## Results Section 2 – Benchmark

### Measuring replay using TDLM and SODA

Before we turn to sequential detection, we briefly summarize the two techniques to quantify replay investigated in this paper, TDLM and SODA. Both methods rely on pattern classifiers to map from neural activity to event probabilities (e.g. viewing an image) for every time point in a time series. If a particular event/class has a (relatively) high probability, given the data, that class is assumed to be “reactivated”, and “replay” is defined as a sequence of such reactivations that follow an expected pattern. How much evidence there is for such an expected sequence of reactivations is often expressed in terms of “sequenceness”, with higher values indicating stronger evidence for replay. One important assumption of both TDLM and SODA is thus that replay events contain activation patterns similar to those on which the classifier was trained during the “localizer” task. While endogenous reactivation does not necessarily have to be identical to sensory evoked activation similar assumptions are routinely made in the context of memory decoding studies.

### Temporally Delayed Linear Modelling (TDLM)

Temporally delayed linear modelling has become popular as a method to quantify sequential spontaneous reactivation using MEG and EEG data. In brief, TDLM first calculates the relation of pairs of time-shifted classifier time courses, for every pair of classifiers, and every time lag of interest with a GLM, similar to a cross-lagged correlation or Granger causality (Granger, 1969). The analysis then checks whether the time-lagged relationships for expected sequence pairs (e.g., A-B) are higher than for other sequence pairs (e.g. F-C), by averaging all beta weights for expected transitions and comparing against the average beta of unexpected transitions. The resulting “sequenceness” score represents the evidence for sequential replay at a specific time lag being present within a time series (in an arbitrary unit scale), with time lag meaning the distance in time between items A and B as well as B and C in the sequence ABC.

TDLM makes several assumptions on the shape and form of the brain activity elicited during replay. First, TDLM assumes that during sequential replay items are reactivated roughly equidistantly in time to each other, with a fixed time lag between each item, that is the same across all participants. Second, TDLM assumes that reactivation peaks are temporally separated and that the temporal resolution of the measurement is at least as high as the replay time lag of interest (i.e., if a time lag of 20 ms is expected, the recording’s sampling frequency must be at least the Nyquist frequency of 100 Hz). Third, at least in its commonly used standard form, TDLM constructs evidence for replay of a full sequence purely based on one-step subsets of transitions (e.g. cumulative evidence for A-> B and B-> C and C - > D), but never verifies the existence of multi-step sequences (A->B->C->D).

### Slope Order Dynamic Analysis (SODA)

SODA was developed for fMRI to address the problem that the sluggish hemodynamic responses and coarse sampling make the detection of fast replay events difficult. The core insight SODA rests on is that largely overlapping classifier responses (of the sort shown in Figure 1D) will still have a predictable pattern of overlap if the underlying events were triggered sequentially. It regresses the probability of all items *during a single* measurement time point to their position in the sequence, asking whether the order of class probabilities corresponds to a hypothesized sequence. Then, the dynamic of the regression slope is inspected over all measurement time points where evidence for sequential activity is assumed. If overlapping, but time lagged sequential activity was present in the signal, the slope is expected to have a bi-phasic shape, i.e., to be negative in an initial period after the neural sequence event, and then positive during a subsequent period (see Figure 1E&F). Magnitude and frequency of this wave like dynamic follow a predictable pattern that depends on sequence speed.

In contrast to TDLM, SODA assumes that the decoded probability increase of individual items is temporally extended and exhibits a clearly overlapping waxing and waning phase related to the haemodynamic response. As long as this assumption is fulfilled, it has fewer constraints on the sampling frequency, allowing for sampling frequencies below 1 Hz to decode neural sequences that occurred at a much faster timescale. SODA uses linear regression to quantify ordering across all transitions, which relaxes constraints on the exact ordering within each replay, i.e., as in any other regression no exact relationship between classification probability and rank is required as long as the overall pattern follows such a relationship. Note, that the choice of regression method is less important, with rank correlation, mean step size between events or linear regression yielding qualitatively similar results, see Wittkuhn & Schuck (2021). Furthermore, SODA finds evidence for entire sequences without looking at individual transitions, as long as the individual reactivations are largely overlapping and especially the first and last position of the sequence are recoverable (statistically speaking, the first and last points have more “leverage” on regression coefficients, see also Figure 6).

### Benchmarking SODA and TDLM on FASTIMAGES

We will now demonstrate how FASTIMAGES can be used to benchmark sequence detection methods by the example of TDLM and SODA. Because the sequences are stimulus-driven and clearly expressed in the neural signal with known onset times, high numbers of repetitions and fewer sources of between trial variance, the dataset provides an upper bound on each method’s sensitivity, representing best-case performance compared to genuine replay studies. Any failure to detect sequences under these conditions would indicate a fundamental limitation of the method. Conversely, the benchmark quantifies the headroom available before a method is applied to the noisier and more ambiguous signals expected during spontaneous replay. We conclude the section by comparing power of TDLM versus SODA via a bootstrap analysis and show effect sizes of detection in this best-case scenario.

### TDLM on MEG data

To decode participant’s brain activity during fast image viewing we trained a logistic regression on the average time point of maximum decoding accuracy (150 milliseconds). For each sequence trial, we estimated the reactivation probability for each image at each time point and normalized the probability time series per trial by dividing by the mean probability per class. We used these probability time series per class as input to TDLM to assess forward and backward sequenceness at any time lag according to the trials’ visual sequence. To normalize sequenceness across participants, we z-scored values per trial across the time axis. The resulting sequenceness curves show clear peaks at the expected time lag per speed condition (Figure 4A).

**Figure 4.**
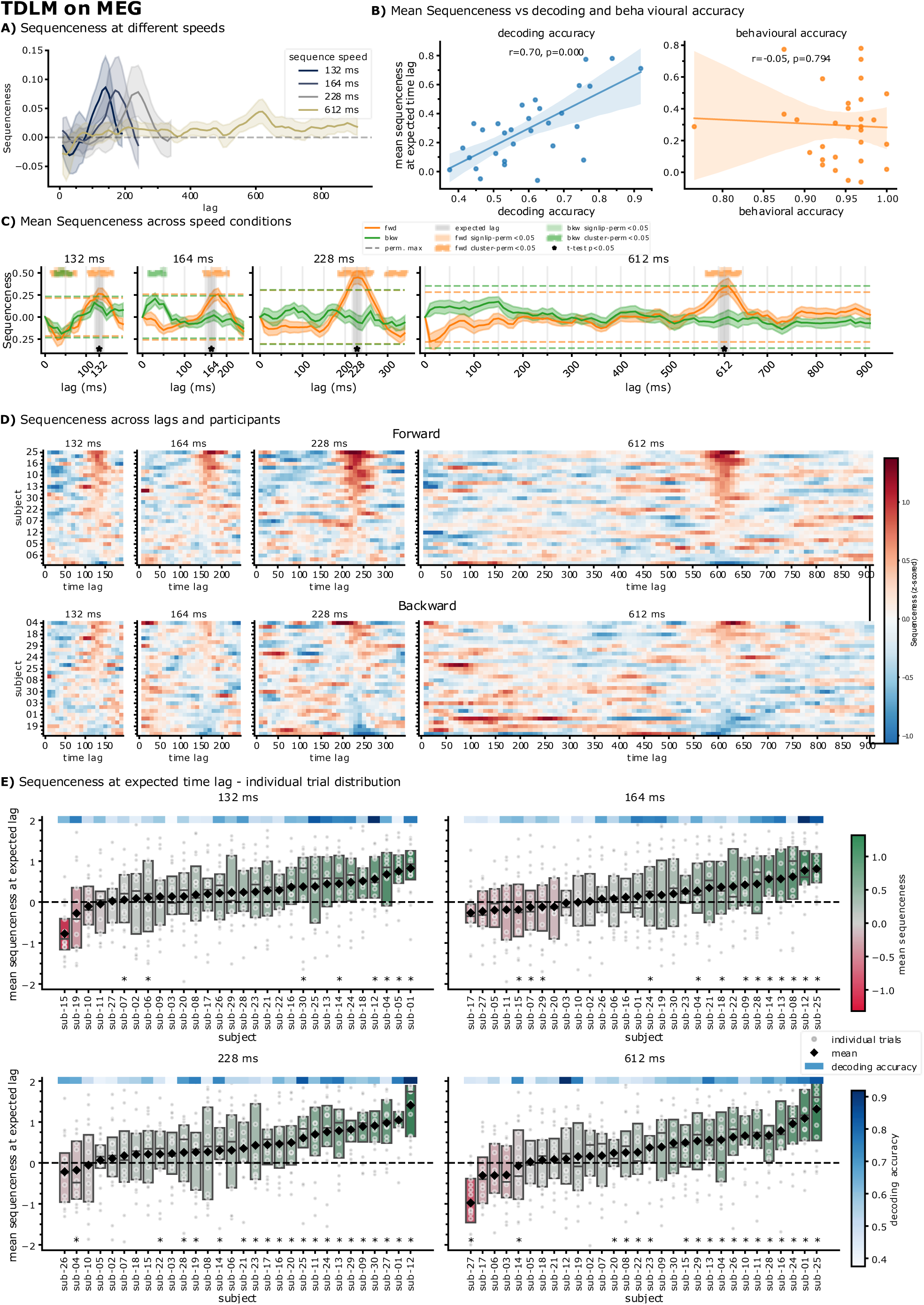
TDLM on MEG – overview of sequenceness results of the sequence trials. A) Mean sequenceness (a.u.) across all four speed conditions in the MEG. Clear peaks are visible at the expected time lag. B) Correlation of mean sequenceness at the expected time lag to decoding performance during the localizer trials (left) and the behavioural accuracy (correct response rate) during the sequence trials (right). Decoding performance was positively correlated with mean sequenceness at the expected time lag in all speed conditions (r=0.7, p<0.001). Right: Mean task performance, as a proxy for attention to the presented stimuli, was not related to mean sequenceness strength at the expected time lag (r=-0.05, p=0.794). C) Average forward and backward sequenceness (z-scored) for the four different speed conditions in the MEG. Three different statistical tests are visualized: A T-test on the mean values at the expected time lag (star), a sign-flip permutation test (darker bands) and a cluster-permutation test (light bands). State-permutation maximum sequenceness values are indicated as a dotted line for legacy purposes. In all conditions, significant sequenceness can be detected at the expected time lag, with the cluster-permutation test finding descriptively larger areas. False positives were present at time range 30-60 ms time lag for the fastest condition. **D)** Mean forward sequenceness per participants. Rows have been sorted by peak sequenceness around the expected time lag (the onset-to-onset interval). To normalize sequenceness between participants, values have been z-scored within each trial. Strong effects are visible in all four speed conditions for the majority of participants. **E)** Overview of sequenceness at the expected time lag per trial and participant. Individual trials are marked with a grey dot. Mean sequenceness per participant is marked with a black diamond. Decoding accuracy during the localizer is shown in a blue band above the plot. Stars signify significance returned by a sign-flip permutation test on trials of that participant. Boxplots are coloured by mean sequenceness from green to red and indicate quartiles and the median. Overall, most participants show a positive effect at the expected time lag. However, mean and median sequenceness across trials is below zero for some participants. Some participants even exhibit negative forward sequenceness in almost all trials. This demonstrates the importance of interpreting TDLM on the group and not individual level.

We averaged the sequenceness for trials of each speed condition per participant and performed a sign-flip permutation test (as proposed in Kern et al., 2026) and a cluster-permutation test on the averaged sequenceness values. TDLM clearly detects the sequences (Figure 4C) and surpasses significance thresholds in all four speed conditions at the expected time lag. False positives were present in the fastest 132-millisecond condition between 30-60 milliseconds that indicated negative forward and backward sequenceness, suggesting that the applied statistics are more likely too liberal than too conservative.

### TDLM’s population variance is explained by decoder performance

While TDLM shows strong effects on the group-level, sequenceness curves show high variance between participants (Figure 4D). While a positive sequenceness at the expected time lag was present in most participants across the sample, this peak was significant using a t-test in only around 50% of participants on average, with only 25% being significant in the fastest, 132 ms speed condition (164 ms: 43%, 228 ms: 64%, 612 ms: 60%). This indicates that even in a best-case scenario, the high variance does not allow conclusions on an individual participant basis. Additionally, when comparing with the group-level plots in Figure 4D, it becomes apparent that some group-level effects are driven by few participants, as is the case for the negative forward peak at 30 ms in the 612-millisecond condition, which was only present in two participants.

Figure 4C shows the variance of sequenceness at the expected time lag per trial, per participant. In all speed conditions, at least some participants exhibit an average negative sequenceness, with some having almost all trials with negative sequenceness at the expected lag. When assessing significance per participant’s trials, only around 50% of participants show significant sequenceness at the expected lag across all speed conditions (132 ms: 26%, 164 ms: 43%, 228 ms: 64%, 612 ms: 60%), highlighting the high noise level per trial. However, per participant only 16 trials were recorded per speed condition, making this analysis slightly underpowered.

Additionally, we assessed whether variance in sequenceness was explained by differences or changes in participants’ behaviour. There was no correlation between task performance (e.g. “Which sequence position was the shoe?”) and mean sequenceness per participant (Figure 4B right). Also, sequenceness was also not related to reaction times of selecting the correct sequence item (Supplement Figure 8). In contrast, decoding accuracy during the localizer was highly significantly correlated with sequenceness at the expected time lag across all speed conditions (*r* =.70, Figure 4B left). This means that the more accurate the decoder was for an individual participant the more sequenceness was detected at the expected time lag.

### SODA on fMRI data

Next, we assessed how SODA performs on the fMRI data. Here, we used the precomputed probability time series provided by Wittkuhn & Schuck (2021), who trained per-class logistic regression classifiers on the mean best decoding time point during the localizer image presentation (TR 4 to 6). We normalized by dividing by the mean probability per class per trial and then applied SODA by regressing the probabilities per TR onto the target sequence. This resulted in one time series of thirteen slopes per sequence trial, per participant, one for each TR. Figure 5A shows a visualization of the slope dynamics over time. Each speed condition shows the characteristic onset (negative slope) and offset (positive) period of the slopes. Note that, per convention, we flipped SODA slopes by-1 to align with the previous paper.

**Figure 5.**
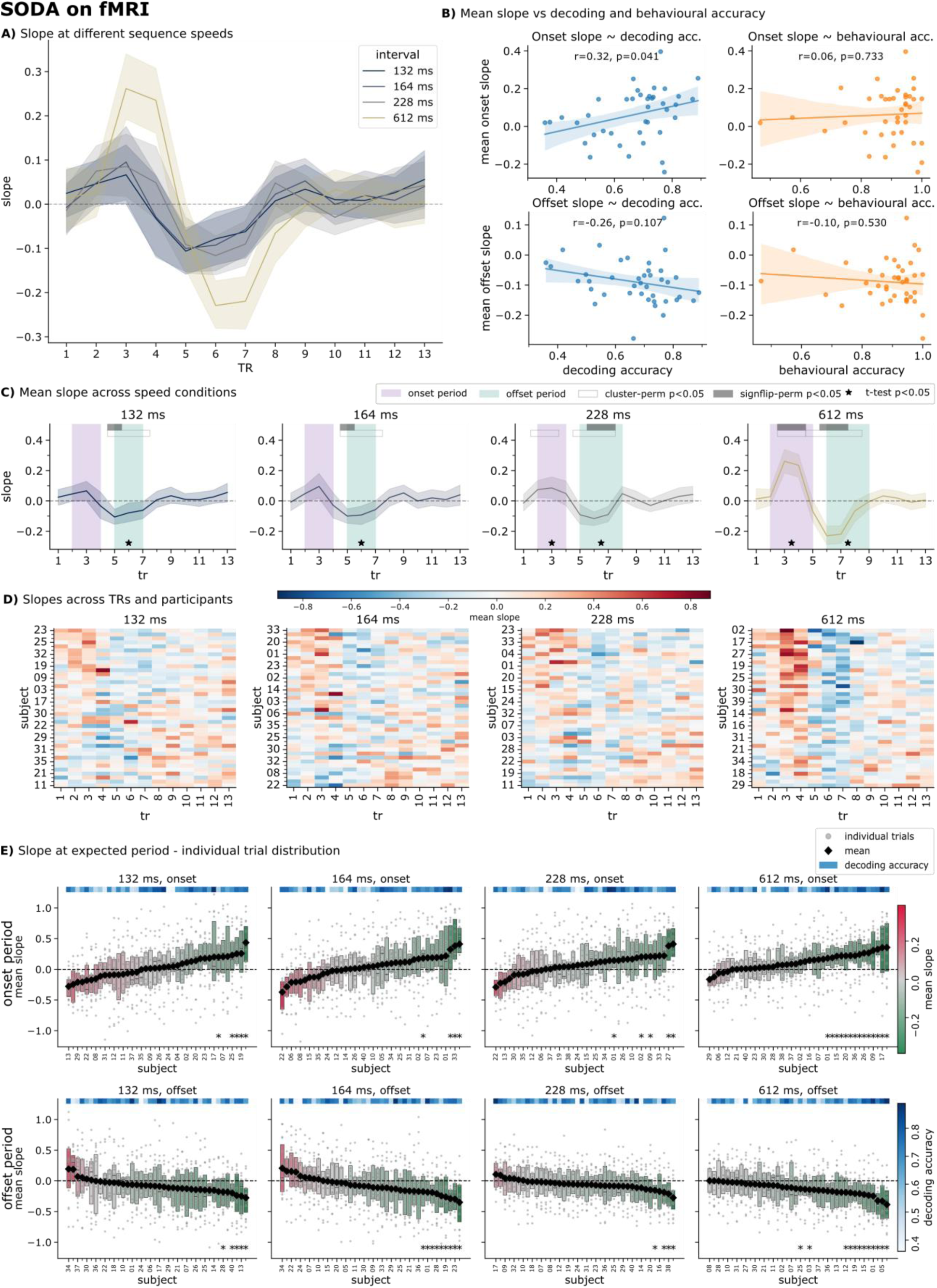
SODA on fMRI – slopes during the sequence trials. **A)** Regression slope coefficients for the different speed conditions across time after sequence onset. Each TR was 1.25 seconds long. **B)** Correlation of decoding accuracy and behaviour with onset and offset mean slope. Left: Decoding accuracy was significantly correlated with the onset slope (upper) but not the offset slope (lower). Right: No correlation between mean behavioural accuracy, as a proxy for attention within the trial, and mean onset or offset slopes. **C)** Mean slope after first image onset across speed conditions across TRs. Onset and offset periods were derived from data of the localizer (see Wittkuhn et al 2021 Table 1). We used three different ways to test for significance: A sign flip-permutation test with t-max correction, a cluster-permutation test and a t-test on averages per period. Slopes were significantly above zero for the offset period in all speed conditions, but less expressed during the onset period, where none of the tests were significant for the fastest two conditions. **D)** Mean slopes per participant across TRs. Rows are sorted by mean slope during the onset period. **E)** Mean slope for onset and offset periods per participant’s trials. Individual trials are marked with a grey dot. Mean slopes per participant are marked with a black diamond. Decoding accuracy during the localizer is shown in a blue band above the plot. Stars signify significance at that time lag as returned by a sign-flip permutation test. Boxplots are coloured by mean slope from green to red and indicate quartiles and the median. Overall, most participants show a positive effect during the onset or offset period. However, mean and median slopes across trials is for some participants below zero. Some participants even exhibit slopes in the opposite direction than expected in almost all trials.

**Figure 6.**
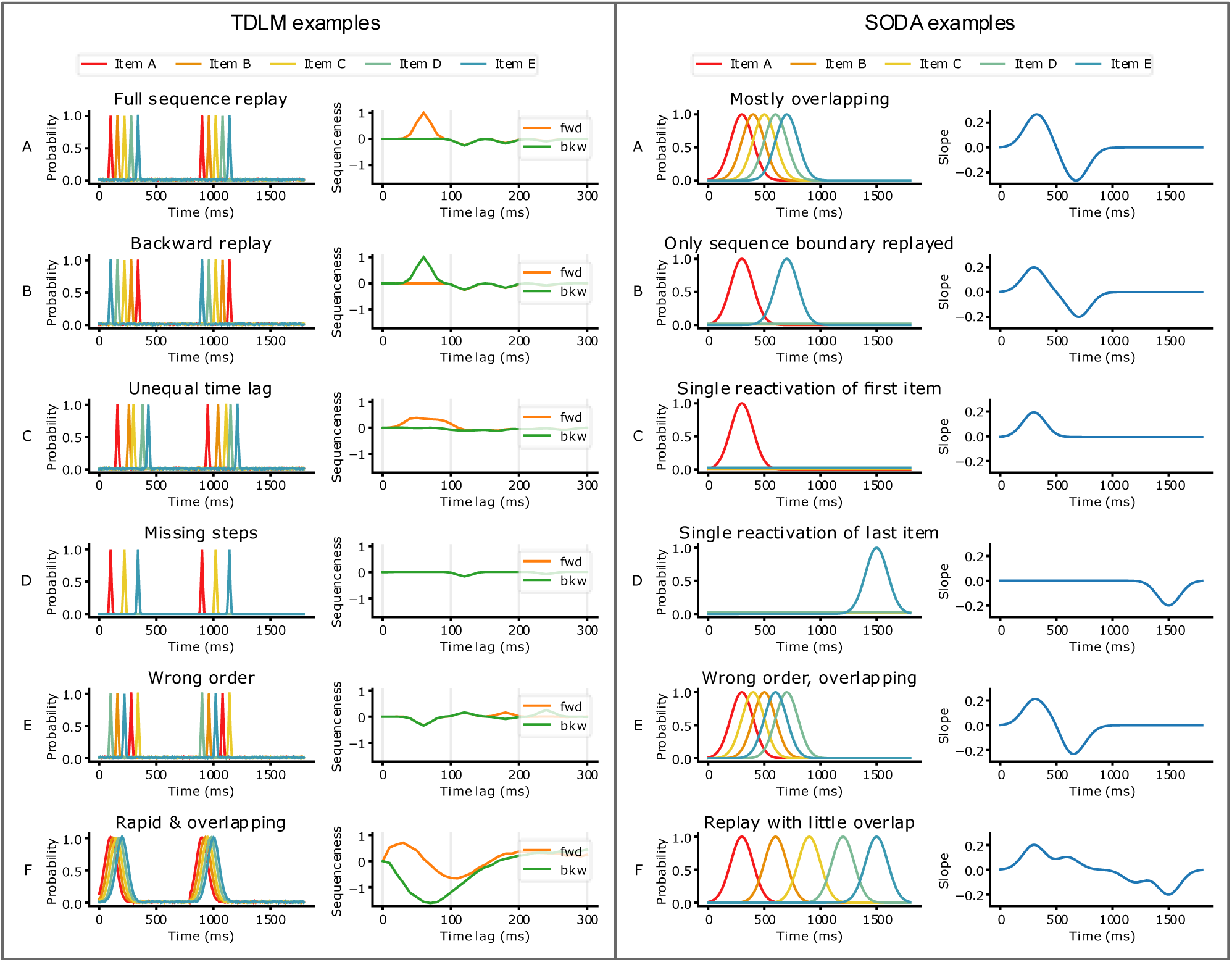
Examples of SODA and TDLM behaviour under different contrived conditions. **Left**: TDLM Examples. **A)** When items are reactivated with constant inter-item lag, TDLM produces a sharp forward sequenceness peak at that lag. **B)** Same as A) but for the backward replay case. **C)** If the inter-item lag varies between transitions, no single lag is consistently expressed and the forward peak smears across multiple lags with reduced amplitude. **D)** When items of the sequence are missing both forward and backward sequenceness stay near chance level. **E)** If the sequence has the wrong order, no clear sequenceness peak is observed. **F)** When reactivations are very rapid and largely overlapping, but sample rate is high, the sequenceness curve expresses no clear peak corresponding to the simulated time lag. See also Appendix of Kern et al. (2026) for more examples. **Right**: SODA examples. **A)** If reactivations are largely overlapping, SODA produces a pronounced onset and offset period. **B)** If only the beginning and end of the sequence are replayed, the resulting slope is almost indistinguishable from a full sequence replay. **C)** A single reactivation of the first item will result in a singular onset period. **D)** A single reactivation of the last item will produce a singular offset period. E) When individual reactivations within the sequence are mixed up, the onset and offset period is still observable. **F)** If there is little overlap, the superposition of single reactivations of the first and last item dominate the slope dynamic.

### SODA on group level

Figure 5C shows the slopes separated by interval speed, together with the predefined onset and offset period, derived from the localizer response function (see Wittkuhn & Schuck, 2021, Table 1). On average, a strong effect is visible for the 612-millisecond condition, with smaller effects visually present for the faster conditions. To assess significance, we performed a sign-flip permutation test with t-max correction and a cluster-permutation test against zero across TRs. For legacy reasons, we also performed a one-sample t-test on the average slopes durin00 the two periods as in Wittkuhn & Schuck (2021). Test results show that the offset period is more sef56nsitive to detection (Figure 5C), being significant in all speed conditions and for both tests. However, SODA struggles to detect the onset period in the faster conditions. The cluster-permutation test was slightly more sensitive and detected the onset period additionally in the 228 ms condition. Furthermore, it became apparent that the onset and offset periods, derived from theoretical models (shaded in red and green) seem to be suboptimal, as they only partially overlap with the empirical clusters revealed by a cluster-permutation test.

### SODA on participant-level

To visualize slope dynamics across the population, Figure 5D shows the mean slopes across the different speed conditions. While slope dynamics can be observed by eye in almost all participants for the slowest condition (612 ms), results were less consistent for the faster conditions with some participants showing no apparent slope mean at the expected times.

Testing for onset and offset periods per participant individually revealed that in the slowest (612 ms) condition more than 80% of participants reached significance while in the faster condition, gradually fewer participants reached significance for both slopes (Figure 5E). However, as with the MEG data, per participant only 15 trials were recorded per speed condition, making this analysis slightly underpowered. Figure 5E shows a boxplot for mean slopes during the onset (upper) and offset (lower) period. Generally, faster speed conditions, on average, have fewer trials recorded with slopes in the expected direction.

Correspondingly, we asked whether attention to the stimuli would explain slope magnitude. However, no correlation between mean onset or offset slope and behavioural performance, as a proxy of attention, was seen (Figure 5B right). However, same as in the MEG, decoder performance during the localizer was positively correlated to the mean slope across all speed conditions, albeit less strongly than in the MEG and only for the onset slope (*r* =.26 and.32; Figure 5B left).

### Transfer of SODA and TDLM to MEG and fMRI

As mentioned previously, SODA was specifically developed for slow and overlapping probability time courses, mainly present in fMRI BOLD signals, while TDLM was developed for the rapid signal changes occurring in M/EEG. In this section, we explore what challenges arise when applying the methods to their non-native field (TDLM to fMRI and SODA to MEG data) and propose adaptations and scenarios leveraging their respective strengths and assumptions. Additionally, we provide a non-exhaustive set of examples that illustrate how the two algorithms behave under different contrived examples that partially violate their assumptions (Figure 6).

### SODA on MEG data

SODA detects whether ordering of probabilities *within* a specific time point adheres to a specific sequential pattern. This necessarily requires a (partial) overlap of the decoded reactivations to be present. In contrast, the decoded activity measured in the MEG is usually rather short-lived. Therefore, in the case of our benchmark dataset (cf. Figure 3C) and mirrored by the results of previous papers showing decoded reactivations to be 10-30 ms in duration (Liu et al., 2019; Wimmer et al., 2020) an overlap of reactivations is mainly present between neighbouring items.

Applying SODA to MEG data (Figure 7) reveals onset slope peaks but no offset phase, because reactivation curves decay too fast for second-item peaks to overlap with the first item’s offset window (see explainer in Figure 6 right). Extending the reactivation period, e.g., by down-sampling the data to a lower sampling frequency or applying a smoothing kernel to the data, or alternatively, creating a classifier that generalizes more in the temporal dimension (see Supplement Figure 5 for our attempt at that) could help.

**Figure 7.**
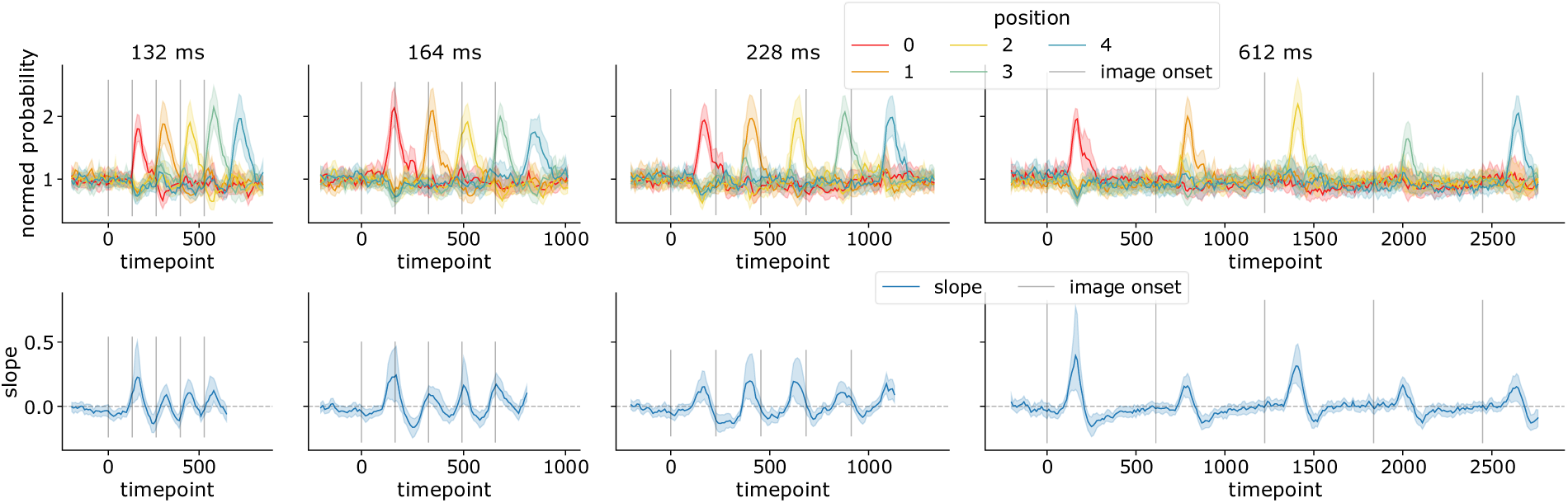
SODA when applied to the MEG sequence presentation. Upper: normalized probability estimates per position (same as Figure 3C). Lower: Mean SODA slope dynamics. While there are offset periods observable, they are due to singular strong activations and not due to sequence replay. Weak overlap between presentations slightly increases offset periods. See Figure 6 right panel for a visual explainer of this phenomenon.

However, previous papers have reported “clustered” reactivation, in which items are reactivated seemingly simultaneously (Kern et al., 2024; Wimmer et al., 2020). Clustered replay is indistinguishable from very fast replay below the sampling rate of the signal, a context in which SODA excels. Therefore, SODA is an excellent candidate to investigate this fast replay in MEG data.

### TDLM on fMRI data

TDLM expects brief, temporally separate reactivation events, a property that fMRI BOLD fundamentally violates. While in principle, TDLM can be applied to fMRI data (as demonstrated in Huang et al., 2024), its ability to uncover faster replay-like events remains severely limited. With a typical TR of 1.25 s (0.8 Hz) the minimum recoverable time lags are multiples of 1.25 seconds, limiting it to slow sequences of reactivations, contrary to expected replay speed. Additionally, in a non-event-driven design, TRs unevenly sample the temporal space, further complicating the TDLM assumptions. Furthermore, individual fMRI trials have a lower signal-to-noise ratio than MEG trials. Applied to the benchmark no speed condition is recoverable (Figure 8 left). Wittkuhn et al. (2021) additionally recorded a control condition with an onset-to-onset interval of 2148 milliseconds, which should be recoverable. However, individual fMRI signal-to-noise of trials is too low, such that significance was only reached when applying TDLM to averaged trials (Figure 8 right).

**Figure 8.**
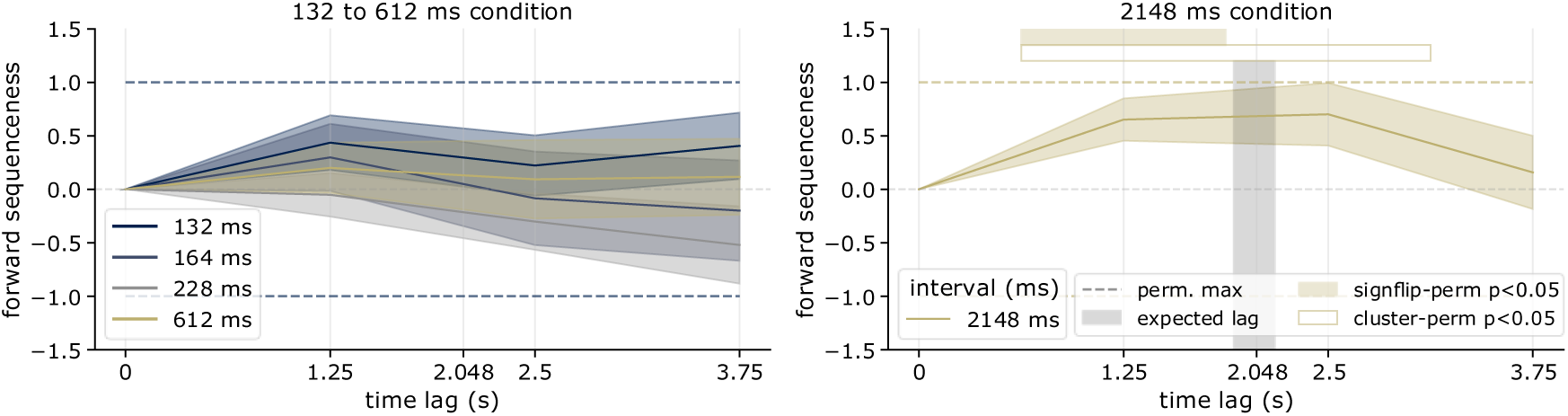
TDLM on fMRI, including a condition with an onset-to-onset interval of 2148 milliseconds. Left: In the sequence conditions in which sampling rate (TR=1.25) was slower than presentation speed, no sequenceness can be uncovered. Right: In a condition in which images were shown very slowly with an onset-to-onset interval of 2148 milliseconds, sequenceness is significantly above zero (tested with a sign-flip permutation test and a cluster-permutation test).

Two options are conceivable: First curve-fitting the sparse TR samples, trying to uncover the original BOLD dynamics. However, measurements are probably too noisy and averaging trials when replay onsets are unknown is not possible. Second, decreasing the TR. However, reducing the TR to 650 milliseconds in an unpublished study, the signal-to-noise ratio decayed dramatically, making decoding worse, not better. In summary, TDLM on fMRI is possible in principle but practically limited to slow sequential processes. However, advances in field strength or acquisition may eventually close the gap.

### Effect size and power of TDLM and SODA

To assess the magnitude of effects measured by TDLM and SODA, we calculated between-participant Cohen’s d effect sizes (Figure 9A) on mean sequenceness and slopes at the expected time points. In addition, to arrive at a single measure per method, for SODA we combined slopes by taking and adding the means per slope and multiplying by 0.5 (onset period) and-0.5 (offset period) and using a t-test to compare against zero. TDLM demonstrated relatively stable, mean medium effect sizes across the speed conditions of d = 0.77 ranging from d = 0.52 (164 ms) to d = 1.12 (228 ms). In contrast, SODA effect sizes were speed-and period-dependent. Mean onset period effect size was d = 0.43 and was markedly reduced at faster interval speeds (d = 0.16 at 132 ms to d = 0.88 at 612 ms), whereas offset period magnitudes (mean d = 0.9) remained stable for fast conditions and increased significantly for slower intervals (d = 0.64 at 132 ms to d = 1.36 at 612 ms). Overall, TDLM effect sizes could not be shown to differ from SODA effect sizes for the combined period measure (independent Z on Cohen’s d, 132 ms: z=0.68, p=0.497; 164 ms: z=0.01, p=0.991; 228 ms: z=1.09, p=0.277; 612 ms: z=-0.7, p=0.063).

**Figure 9.**
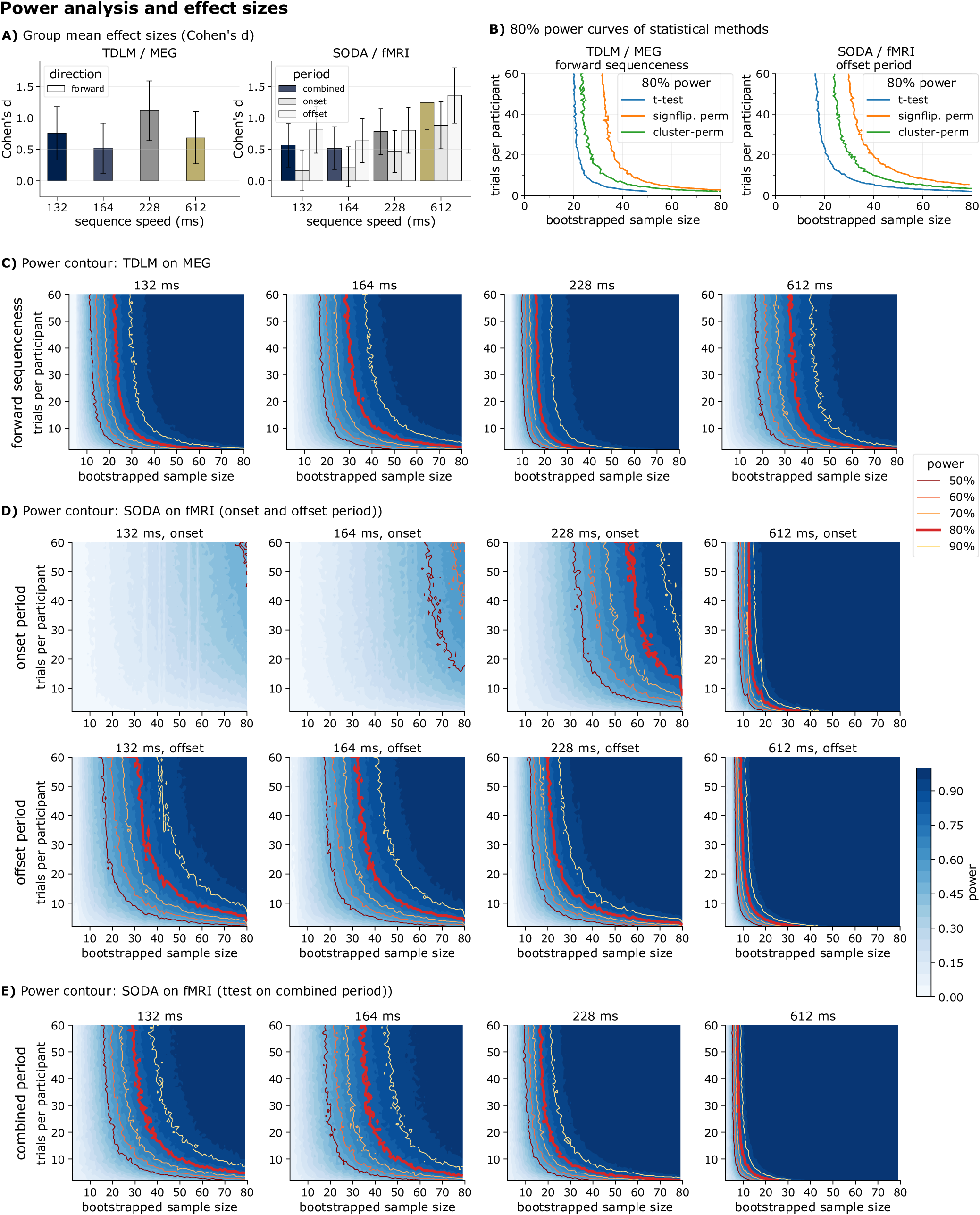
Power analysis and effect sizes for TDLM on MEG and SODA on fMRI data **A)** Between-participant Cohen’s d effect size and 95% confidence intervals for the different speed conditions. Left: Effect size of TDLM mean sequenceness at the expected time lag (+-10 ms) in the MEG. Right: Effect size of SODA mean slope during the predefined onset and offset periods and a combined slope measure. **B)** 80% power for TDLM and SODA in their respective modality. We bootstrapped number of participants and trials per participant until an average power of 80% was reached for each statistical method. Left: 80% power lines for TDLM and MEG on the average forward sequenceness at the expected time lag. Right: 80% power lines for SODA and fMRI on the average offset period slope. For visualization purposes we only display the offset periods, as faster onset periods in the fMRI do not reach 80%. **C)** TDLM power contour plot for bootstrapped samples of trials and participants for the four interval speeds. For each combination of sample size and trial number, 1000 draws of participants were resampled with replacement, and for each draw, trials were resampled with replacement per participant. The three before-mentioned statistical tests were applied and checked for significance. The resulting contour plot shows mean power of the three methods as a function of bootstrapped sample size and number of trials, with the bold red line indicating the threshold for 80% power. For plots separated by statistical test, see Supplement Figure 9 and Supplement Figure 10. **D)** same as C) but for SODA. **E)** Power contour as in D but for a combined onset and offset period by taking the mean of the average onset and negative offset period.

To assess the power of TDLM and SODA to detect an effect with 80% chance, we performed a bootstrap analysis across increasing numbers of simulated trials and participants. For each combination of sample size and trial number, we chose a list of participants with replacement and resampled the trials per participant with replacement, repeating the procedure 1000 times. We tested for significance using all three methods as defined above and averaged their values. For the methods without assumptions of the time point of effect, we counted a successful detection if curves were significantly above zero within the expected time point or lag (onset or offset period for SODA and expected time lag +-10 ms for TDLM). The resulting power contour plot in Figure 9C&D, as proposed in Baker et al. (2021), shows power for each combination of bootstrapped participants and trials per participant.

For TDLM, power is stable across all speed conditions, with the strongest power being in the 228 ms condition. Generally, for both methods, power drops off significantly with fewer than 15 trials (Figure 9C). For SODA it becomes apparent that the onset period in the fastest two conditions (132 and 164 ms) is not suitable to detect an effect with confidence with any of the simulated sample and trial sizes when using separate onset and offset periods (Figure 9D upper row left). Furthermore, SODA’s power decreased with increasing interval speeds. Nevertheless, when using the combined measure as defined above, SODA’s power was markedly increased (Figure 9E).

Lastly, we wanted to compare the power of the three different statistical methods used in the context of TDLM and SODA: t-test on averaged values during the expected time bin or lag, sign-flip permutation testing with t-max correction and cluster-permutation testing. Figure 9B shows the 80% power line for the three statistical test methods. For both TDLM and SODA, the t-test on the mean values at the expected time point had the highest power. However, when expected time lags or bins are not known a priori, the cluster-permutation test was superior to the sign-flip permutation test. Note, while this shows the power to detect an effect, it does not indicate the false positive rate, which we did not examine in this context.

## Discussion

In this paper, we introduce FASTIMAGES, a combined MEG and fMRI dataset containing fast neural sequences of known structure, designed to validate and compare neural sequence replay detection methods in human neuroimaging data. Using this dataset, we benchmarked two algorithms for human replay detection, TDLM and SODA, and assessed their sensitivity, effect size and statistical power through a bootstrap analysis. We show that both algorithms perform well in their designated target modality (i.e., MEG for TDLM and fMRI for SODA) while transfer to the non-target modality proved challenging. We show that in both modalities sequence detection significantly depended on the localizer data and quality of the decoder.

TDLM detects sequenceness in the MEG data in all four speed conditions (132, 164, 228 and 612 ms onset-to-onset interval) and clearly surpasses significance thresholds at the expected time lag while showing effect sizes (Cohen’s d) in the medium range. SODA evidence for sequenceness contains two signatures, measured during an onset and an offset period. SODA showed significant effects in the offset periods across all speed conditions with comparable effect sizes to TDLM as applied in MEG. Detection in the onset periods was much more difficult, and only expressed significant effects for slower speeds (228 and 612 ms). An integrated measure, that combines onset and offset periods, showed markedly increased power and effect sizes. However, it is only applicable in event-related paradigms, as onset and offset periods must be known a priori. For SODA, effect sizes decreased with faster speeds, consistent with the increasing overlap of BOLD responses of the individual stimuli. On average, TDLM and SODA exhibited similar power to detect an effect with 80% chance with around 20 participants and 20 trials necessary when expected time lags of interest, or onset and offset periods are known beforehand. Nevertheless, power was stable across all conditions for TDLM, while SODA had higher power in slower (228 & 612 ms) and lower power in faster conditions (132 & 164 ms). A comparison of three different ways to test for significance revealed that when time points for time lags and slope periods are unknown that cluster permutation testing (as in fMRI) is consistently more powerful than a sign-flip permutation test. Hence, we recommend using cluster permutation tests as best-practice. Of note, TDLM and SODA are most frequently used for detection of replay events for which the exact timing is unknown (Liu et al., 2019, 2022; Wimmer et al., 2020), in our benchmark we did not take into account the search space, which likely will increase the required sample size and trial number required (Kern et al., 2026). For TDLM, the search space could be furthermore reduced by deriving an expected lag from the literature (Buzsáki, 2015; Fernández-Ruiz et al., 2019).

A strength of TDLM is that it offers a clear and straightforward interpretation, yielding an exact time lag at which there is evidence for a sequence of reactivations. The resulting sequenceness curve can be tested for significance using various methods. Furthermore, the GLM backbone makes TDLM flexible to incorporate covariates and controls that can be inserted into the design matrix (as is often done with alpha-oscillatory control, e.g., Liu, Dolan, et al., 2021; Liu et al., 2019; Wimmer et al., 2020). Beta values within the GLM are interpretable and traceable to certain transitions and time lags enabling mechanistic examinations (Agresti, 2015; Liu, Dolan, et al., 2021; Nelder & Wedderburn, 1972). Additionally, although sensitivity of TDLM benefits from data being event-related, it is in principle invariant to the temporal distribution of replay onsets within a measurement period, allowing for larger search spaces, e.g., in resting state. However, the size of the search space may require unrealistically high rates of replay (Kern et al., 2026). This issue may be solved by restricting analyses to certain M/EEG microstates (Michel & Koenig, 2018; Xu et al., 2020) or other phenomena known to be associated with replay such as theta waves (Burke et al., 2014; Fuentemilla et al., 2010), sleep spindles (Abdellahi et al., 2024; Pedrosa et al., 2024), or slow oscillations (Cousins et al., 2014; Genzel & Robertson, 2015; Rasch et al., 2007).

TDLM has several implicit assumptions and prerequisites, including an appropriately high sampling rate. It assumes that peaks of sequences of reactivations are temporally separate and that replay speeds remain relatively stable within and between participants. In the case of the benchmark this stability is built into the data set by presenting images with a fixed onset-to-onset interval. Given the spread of individual peak frequencies of canonical oscillations in other domains (Buzsáki & Draguhn, 2004; Haegens et al., 2014; Mahjoory et al., 2020) and previous TDLM studies reporting a wide range of time lags (from 30 to 150 milliseconds, e.g., Eldar et al., 2020; Kurth-Nelson et al., 2016; Liu et al., 2019; Wimmer et al., 2020), it is unclear if all participants would replay sequences at strictly the same speed. Additionally, while multi-step TDLM has been proposed (Liu, Dolan, et al., 2021), the majority of studies utilize cumulative one-step transition evidence to quantify replay. Therefore, in principle, when testing for A->B->C->D, neural data of several strong A->B transitions could reach significance and would be indistinguishable from full-sequence replay without further analysis. Finally, if replay occurs faster than the typically used sample rate’s Nyquist frequency of 50 Hz, TDLM would be unable to detect it as clear peaks are missing in this case.

We also applied TDLM to fMRI data, where it would not be expected to work in light of these assumptions. Hence, TDLM on fMRI has proven difficult and previous studies failed to find effects as well (Huang et al., 2024). Without clear probability peaks and sparse sampling, TDLM cannot infer any temporal dependency. While workarounds such as fitting response curves on TR-spaced data (Cohen, 1997; Miezin et al., 2000) are in principle conceivable, we expect them to perform poorly given the low SNR of single-trial fMRI. Crucially, TDLM works well when expected time lags are above the TR. As shown, it successfully uncovers sequenceness in the 2148 ms condition. Therefore, paradigms in which sequential brain activity is expected over a longer time frame may be a promising target of fMRI-TDLM, for example during episodic recall (J. Chen et al., 2017) or behavioural sequences (Tang et al., 2021). Nevertheless, it is unclear which processes exist on that time scale that also fulfil TDLM’s assumption of fixed time lag across the paradigm and population. In essence, the benchmark we provide allows testing sensitivity of algorithms relying on fast non-overlapping sequences at high sampling rates and showing that their utility breaks down for data of a different structure. It is of course possible that algorithms can be developed that are able to bridge the two worlds, for which the benchmark again provides the ideal test bed.

SODA has been developed to overcome the limitations of fMRI in the context of replay detection. It detects sequentiality in data with low sampling rates and slow and largely overlapping readouts of brain activity. In this case, a characteristic onset and offset period is expected to reflect the changing slope dynamic of a regression of the reactivation probabilities per time point on the sequence order. SODA regresses onto the entire sequence at once, yielding evidence for the full sequence that is more robust against exact replay time lags and replay speeds. Speed itself can be estimated via a frequency spectrum analysis on the slope dynamic, albeit with less fidelity than with TDLM (Wittkuhn & Schuck, 2021). SODA in its current version has mostly been applied to event-related paradigms and hence under the assumption that replay onsets are roughly aligned between participants, however extensions using slope frequency analysis have been proposed (Wittkuhn & Schuck, 2021).

However, a major caveat for the interpretation of SODA results arises from its reliance on the onset and offset period. First of all, the theoretically derived periods only partially overlap with the empirical clusters (see Figure 5C). Second, interpretation becomes tricky if only one of the two periods is significant, as is the case in the faster conditions (132 and 164 ms). This leads to the complication of replay directionality: If only one clear period can be observed, it is impossible to say whether it is the onset phase of a forward replay (Diba & Buzsáki, 2007) or the offset phase of reverse replay (Foster & Wilson, 2006). While this is partially solved by combining the onset and offset phase into a combined measure this is only applicable if onset and offset period timings are clearly known, as is only the case during event-related paradigms. In this case, a model fit to an expected decoding response curve might also be applicable. As Wittkuhn & Schuck (2021) mention, the effects in their analysis were driven by the first and last item of the sequence (see Figure 6 right panel). On the one hand, this makes SODA resilient to outliers, out-of-sequence replay or missing reactivation events within the sequence. On the other hand, any single reactivation will produce a slight offset phase if its class is at the start of the regressed sequence (and vice versa an onset phase if the last item of the expected sequence is reactivated individually), as long as other probabilities remain low during the same time period (Figure 6 right, B to E). Therefore, further methodological advancements must be made to uncover onset and offset periods separately for replay speeds below 228 milliseconds state-to-state lag in non-event related paradigms.

As proof of concept, we assessed the feasibility of applying SODA to the MEG data. As would be expected, it becomes clear that main assumptions about the source signal are violated and hence SODA is not readily applicable to MEG. Decoded reactivations in the MEG are brief and even in the fastest speed condition only slightly overlapping. Therefore, the distinct change from onset to offset phase of the slope dynamic is missing. While signals of FASTIMAGES could potentially be spread temporally, e.g., by Gaussian smoothing or changes in the classifier, a smoothing would also reduce the signal strength and attenuate potential brief reactivation bursts during endogenous replay.

Lastly, we want to highlight one potential use case of SODA in M/EEG. Previous research has found clustered reactivation, in which several stimuli are seemingly reactivated simultaneously (Kern et al., 2024; Wimmer et al., 2020). However, due to TDLM’s nature, such clustered replay would be indistinguishable from rapid replay faster than the sampling frequency. The necessary overlap for SODA would occur for very rapid replay even in the context of MEG. This could then be used to find out whether clustered reactivation is fast replay in disguise. This is especially promising when one considers that TDLM has previously reported time lags of replay between 50 ms and 200 ms that seem too long for ripple related replay (Buzsáki, 2015).

Both methods exhibit a significant relationship between the sequenceness value and the decoding accuracy of the localizer task, which could not be explained by participants’ attention to the task (measured via attention checks and behavioural performance). It is a mathematical necessity that a better decoding accuracy translates to an increased fidelity of reactivation detection. This is because decoding accuracy limits how much evidence can be accumulated from the endogenous signal, i.e., on average, a decoder with 33% accuracy will only correctly detect 1 in 3 reactivations. More broadly, this means that optimized decoder training and parameter choice are required ingredients for high sensitivity in both methods. In our benchmark, for comparability, we deliberately kept our decoder training parameters equivalent to previous publications (Kern et al., 2024; Wittkuhn & Schuck, 2021). As most studies do, we trained decoders on the time point of maximum average decoding accuracy across the sample, yet individual participants’ peaks varied considerably (Supplement Figure 3). A classifier trained at a suboptimal time point for a given participant will capture a suboptimal neural representation, directly reducing the sensitivity of downstream sequence detection. As has been proposed elsewhere, improved measurement precision can strongly benefit resource intense studies in human neuroscience (Nebe et al., 2023). Hence, future studies should systematically evaluate how decoder training choices affect the sensitivity and specificity of sequence detection methods, for instance by comparing classifiers trained at individual versus group-level optimal parameters, by varying the training time window, or by exploring alternative decoding architectures altogether. Such parameter sweeps are readily feasible with the FASTIMAGES dataset and would help disentangle methodological sensitivity from true neural effects. Ideally, future studies will themselves collect ground truth data with the paradigm used in FASTIMAGES, thereby allowing authors to demonstrate the sensitivity of their implementation of replay detection. Nevertheless, it remains an open question whether decoders trained on stimulus-evoked activity during a localizer task transfer well to the internally generated reactivations, such as those occurring during spontaneous replay (Bezsudnova et al., 2024; Vikbladh et al., 2024). Localizer-trained classifiers capture sensory-driven activation patterns, which may differ in spatial distribution, temporal profile, or signal strength from endogenous reactivations (Dijkstra et al., 2021).

Finally, future research should leverage the FASTIMAGES dataset to move beyond validation and towards systematic methodological optimisation. First, the partial mismatch between theoretically derived and empirically optimal onset/offset windows for SODA, together with TDLM’s assumption of fixed time lags, motivates the development of more flexible models that accommodate variable replay speeds, data-driven window definitions, and robustness to partial or non-contiguous sequences, all of which can be tested and calibrated using FASTIMAGES. Second, the complementary strengths of TDLM and SODA invite hybrid approaches, e.g., by formulating SODA as a GLM and performing computations, as in TDLM, as a combination of one-step transitions might reduce the necessity for overlap of the entire sequence. At an abstract level, SODA regresses the sequence onto each time step, while TDLM regresses the sequence on row-wise time shifted versions of the probability vectors and then sums evidence for expected transitions. These similarities should be further explored and expanded.

### Limitations

While FASTIMAGES is intentionally framed as a best-case scenario dataset, several limitations need to be considered when generalizing to neural events in other contexts (e.g. replay). First, FASTIMAGES follows an event-related design, inducing neural sequences with a known time lag by visually presenting images one after the other. Therefore, decoders trained on the exact same stimuli during the localizer task will arguably have higher sensitivity to detect such brain activity than spontaneous non-stimulus-driven brain activity related to the same images. Our dataset therefore likely over-estimates power.

Second, the time-locked nature of the paradigm means that sequence onsets are aligned across trials and participants. Real replay is not time-locked to an external event, which removes the benefit of trial averaging and makes detection considerably harder. Relatedly, due to the nature of visual processing, stimulus-evoked activity during the sequence presentation was spread over a longer time period, while spontaneous reactivations during a replay event are probably much shorter. As SODA is tuned towards longer reactivation, this in turn slightly favoured the paradigm towards SODA.

## Conclusion

FASTIMAGES validates that TDLM and SODA each perform well when the time series data adhere to their requirements. At the same time, FASTIMAGES demonstrates divergence inasmuch as the methods do not transfer readily to modalities that violate these requirements. TDLM requires temporally resolved probability peaks available in MEG but absent in fMRI. SODA depends on overlapping BOLD responses that do not arise in MEG. Hence, the two methods are complementary rather than interchangeable. Across both methods, decoder quality emerged as the dominant limiting factor, opening the possibility of marked improvements when decoders are optimized. Because FASTIMAGES is a best-case scenario with stimulus-driven, time-locked sequences decoded by classifiers trained on the same stimuli, it provides an upper bound on detection sensitivity. We suggest using it as a sanity check for existing pipelines, a testbed for new methods, and a shared reference point for making sequence detection in human neuroimaging more robust and reproducible.

## Acknowledgements

This work was supported by an Emmy-Noether Grant (404679789) and an ERC consolidator grant (101170886) to GBF, a German Academic Scholarship Foundation stipend to SK. NWS was funded by the Federal Ministry of Education and Research (BMBF) and the Free and Hanseatic City of Hamburg under the Excellence Strategy of the Federal Government and the Länder and a Starting Grant from the European Union (ERC-StG-REPLAY-852669).

## Methods

### Participants

Seventy healthy participants took part in the study. Forty young and healthy adults were recruited for the fMRI experiment (age: M = 24.6 years, SD = 3.7 years, range: 20-35 years, 20 female, 20 male). All fMRI participants were right-handed, had corrected-to-normal vision, and spoke German fluently. The fMRI dataset and task are described in detail in Wittkuhn & Schuck, (2021). Thirty additional participants were recorded in the MEG using an adapted version of the same paradigm using the same inclusion criteria (age: M = 24.2 years, SD = 2.9 years, range 20-35 years, 9 male, 21 female). All participants provided written informed consent. The ethics commission of the German Psychological Society (DGPs) approved the fMRI study protocol (NS 012018) and the Ethics Committee II of the University of Heidelberg approved the MEG study protocol (2023-634).

### Experimental design

The five stimuli were grayscale images of a cat, chair, face, house, and shoe (400 x 400 pixels each) taken from Haxby et al., (2001), which have been shown to reliably elicit object-specific neural response patterns in previous studies. The task was programmed in MATLAB using Psychtoolbox (fMRI) and PsychoPy 2023 (MEG). The experiment consisted of two main conditions: slow localizer trials, fast sequence trials. In the fMRI, an additional condition was recorded, which we do not report (repetition trials).

### fMRI task variant

Slow (localizer) trials. Localizer trials were designed to elicit object-specific neural response patterns for classifier training. Five images were presented individually in pseudo-randomized order. Each trial began with a fixation dot (300 ms), followed by one image for 500 ms, and a variable inter-stimulus interval drawn from a truncated exponential distribution (mean 2.5 s, lower limit 1 s). Participants performed an oddball detection task, pressing a button when an image appeared upside-down (20% of trials). Only neural activation patterns from correct rejection trials (upright stimuli, no response) were used for classifier training. Across the fMRI experiment, 120 unique sequences of five images were presented (600 stimulus presentations total, 120 per image category), distributed across 8 functional runs in two sessions. Slow trials were interleaved with sequence and repetition trials.

Fast sequence trials. During sequence trials, participants were shown sequences of all five images at varying presentation speeds. Each trial began with a target cue word (e.g., “shoe”) presented for 1000 ms, followed by a blank screen for 3850 ms to reduce interference between cue-evoked and sequence-evoked neural patterns. A brief fixation dot (300 ms) signalled the onset of the upcoming sequence. All five images were then presented in rapid succession, each for 100 ms, separated by an ISI of 32, 64, 128, 512, or 2048 ms (fMRI). The ISI was constant within a trial. The ground-truth temporal distance between two successive images (onset-to-onset interval) was therefore 100 ms (image duration) plus the ISI.

Following the sequence, participants waited through a delay period (until 16 s from sequence onset had elapsed), during which they listened to bird sounds as a distraction. Participants then reported the serial position of the cued target image by choosing between two response options displayed on screen (1.5 s response window). The serial position of the target was drawn from a Poisson distribution (lambda = 1.9, truncated to 1-5) such that targets appeared more often at later positions, discouraging early disengagement from the sequence. Auditory feedback (coin sound for correct, buzzer for incorrect) was provided after each response.

In the fMRI experiment, 15 unique sequences were selected per participant, chosen such that each image appeared equally often at the first and last position. Each sequence was presented at all 5 ISI speeds, yielding 75 sequence trials total across 8 functional runs in two sessions. Sequences were distributed across 8 groups of participants to ensure all 120 possible orderings were used equally often across the sample. Each sequence speed was paired with a fixed image order for a given participant (i.e., one specific permutation of the five images was always presented at a given ISI). The code for the fMRI task variant, which was programmed using PsychToolbox can be found on GitHub: https://github.com/lnnrtwttkhn/highspeed-task.

### MEG task variant

The MEG task followed the fMRI task conceptually but was slightly adapted: Slow trials were no longer interleaved but a single localizer session was recorded. No repetition trials and no 2048 sequence condition was recorded. Fixation cross and wait times were reduced to accommodate scanning only a single session.

Slow (localizer) trials: In the MEG experiment, 160 localizer trials were recorded during a single session (32 per stimulus). The sequence of localizer images was balanced via a DeBruijn-sequence (DeBruijn, 1946), such that each transition of images was occurring equally often. Eight blocks were performed, each separated by a short break of 30 seconds. In each trial, a fixation cross was shown for 300 ms, then the image for 500 ms after which an ITI fixation cross was shown, drawn from a truncated exponential (mean 2.5 s, lower limit 1 s). In 20% of the cases the image was upside down, which needed to be indicated via a button press.

Fast sequence trials: In the MEG experiment, 16 sequence trials were recorded per speed condition (64 total) in a single session. Eight different sequences of the possible 120 permutations were shown per participant with transitions balanced between participants. Same as in the fMRI, each trial began with a target cue word (e.g., “shoe”) presented for 1000 ms, followed by a blank screen for 1 s. A brief fixation dot (250 ms) signalled the onset of the upcoming sequence. All five images were then presented in rapid succession, each for 100 ms, separated by an ISI of 32, 64, 128, 512. The ISI was constant within a trial.

Following the sequence, participants waited through a delay period (until 8 s from sequence onset had elapsed), during which they listened to bird sounds as a distraction. Participants then reported the serial position of the cued target image by choosing between two response options displayed on screen (2.5 s response window). The serial position of the target was drawn from a Poisson distribution (lambda = 1.9, truncated to 1-5) such that targets appeared more often at later positions, discouraging early disengagement from the sequence. Auditory feedback (coin sound for correct, buzzer for incorrect) was provided after each response. The code for the MEG task variant, which was programmed in PsychoPy, can be found on GitHub: https://github.com/CIMH-Clinical-Psychology/highspeed-task-MEG.

### Study procedure

The fMRI study consisted of two experimental sessions (approximately 2.5 h and 1.5 h, respectively). In the first session, participants provided informed consent, completed a demographic questionnaire and a Digit-Span Test, and performed a 10-min practice of the main task. Subsequently, participants entered the scanner for a 5-min pre-task resting-state scan, four functional task runs of approximately 11 min each, a 5-min post-task resting-state scan, fieldmap acquisitions, and an anatomical scan. The second session was identical except participants entered the scanner immediately. The MEG study consisted of a single session (around 2.5 h), equivalent to the first fMRI session. After the MEG session, an anatomical scan (MPRAGE) was conducted in a 3T Siemens scanner.

### fMRI data acquisition and preprocessing

fMRI data were acquired on a research-dedicated 3T Siemens Magnetom TrioTim scanner with a 32-channel head coil at the Max Planck Institute for Human Development in Berlin, Germany. Functional images were acquired using a T2*-weighted multi-band EPI sequence (64 slices in interleaved ascending order, anterior-to-posterior phase-encoding direction, TR = 1.25 s, TE = 26 ms, voxel size = 2 x 2 x 2 mm, matrix = 96 x 96, FOV = 192 x 192 mm, flip angle = 71 degrees, MB acceleration factor 4). Slices were tilted 15 degrees forward relative to the rostro-caudal axis to improve hippocampal signal quality. Each of the two sessions included four functional task runs of approximately 11 min (530 volumes each) and two 5-min resting-state runs (233 volumes each). High-resolution T1-weighted MPRAGE anatomical images were also acquired (256 slices, TR = 1900 ms, TE = 2.52 ms, voxel size = 1 x 1 x 1 mm).

Preprocessing was performed using fMRIPrep v1.2.2 (Esteban et al.) and included head-motion correction (FSL mcflirt), slice-timing correction (AFNI 3dTshift), co-registration to the T1-weighted reference using boundary-based registration (FreeSurfer bbregister), susceptibility distortion correction using blip-up/blip-down field maps, spatial normalization to MNI152NLin2009cAsym space via nonlinear registration (ANTs), and spatial smoothing with a 4 mm FWHM Gaussian kernel (FSL SUSAN). Brain surface reconstruction was performed using FreeSurfer. fMRI data were detrended separately for each run across all task conditions to remove low-frequency signal drifts. Full details of the preprocessing pipeline are reported in Wittkuhn & Schuck (2021).

### MEG data acquisition and preprocessing

MEG data were recorded using a MEGIN TRIUX system with 306 sensors (102 magnetometers, 204 planar gradiometers) at a sampling rate of 1000 Hz. Five continuous head position indicator (cHPI) coils were applied and their positions digitized together with anatomical landmarks using a Polhemus Fastrack system. Horizontal and vertical EOG as well as ECG were recorded for later artifact detection. T1-weighted anatomical MRI images were acquired for each participant and processed using FreeSurfer’s recon-all for cortical surface reconstruction.

MEG preprocessing was performed using MNE-Python v1.8 (Gramfort, 2013) using MNE-BIDS-Pipeline. Noisy and flat channels were automatically detected using find_bad_channels_maxwell and marked as bad. Missing or broken MEG channels were replaced with zero-filled channels using the correct sensor geometry from a template, marked as bad, and interpolated. Continuous head position was estimated from the cHPI coils (compute_chpi_amplitudes, compute_chpi_locs, compute_head_pos). Signal-space separation with temporal extension (tSSS, Taulu & Simola, 2006) was applied together with continuous head movement compensation (mne.preprocessing.maxwell_filter). Additionally, extended SSS (eSSS) was applied using empty-room projection vectors (3 gradiometer and 3 magnetometer components, computed via mne.compute_proj_raw with meg=“combined”) from the empty-room recording closest in time to each participant’s session (Helle et al., 2021). All runs (main task, two resting-state runs) were transformed to a common head position defined by the average head position during the main task run (compute_average_dev_head_t). Data was down sampled to 100 Hz using MNE’s default settings. Epochs were extracted time-locked to stimulus onsets. Localizer epochs spanned-0.2 to 0.8 s relative to image onset (71 time points at 100 Hz). For individual fast image analysis, epochs spanned-0.2 to 0.5 s relative to each image onset within the fast sequence. For TDLM analysis, full sequence epochs were extracted spanning the entire sequence duration plus a buffer to accommodate the maximum tested time lag.

### MEG classifier training

To decode stimulus identity from MEG sensor patterns, we trained L1-regularized logistic regression classifiers in a one-vs-rest scheme (LogisticRegression with L1 penalty, liblinear solver, max 1000 iterations; scikit-learn, Pedregosa et al., 2011). To identify the optimal regularization strength, we performed a grid search over 30 values of the inverse regularization parameter C (ranging from 0.1 to 50) using stratified leave-one-out cross-validation on localizer trials. For each participant, cross-validated decoding accuracy was computed at each time point within the localizer epoch (-0.2 to 0.8 s). The C value yielding the highest mean peak decoding accuracy, averaged across all participants, was selected for subsequent analyses (C=9.1), see Supplement Figure 4.

Final classifiers were trained for each participant on all localizer trials using the group-average best C (9.1) value and the group-average time point of peak decoding accuracy (150 ms post-stimulus). These classifiers were then applied to the sequence trials to obtain stimulus-specific reactivation probabilities at each time point.

### fMRI classifier training

For the fMRI data, we used the precomputed classification probabilities provided by Wittkuhn & Schuck (2021). In brief, an ensemble of five independent binary classifiers (one per stimulus class) was trained using multinomial logistic regression with L2 regularization (C = 1.0, lbfgs solver, max 4000 iterations; scikit-learn v0.20.3) in a leave-one-run-out cross-validation scheme (8 folds). For each class-specific classifier, all other classes were relabelled to a common “other” category, and class weights were adjusted inversely proportional to class frequencies. Training used only correct rejection trials (upright stimuli, no response) from the slow/localizer condition.

Feature selection combined a functional ROI approach with anatomical masks. For each cross-validation fold, a first-level GLM (SPM12) was fitted to the training data, modelling stimulus onsets as boxcar functions (500 ms duration). The resulting t-maps were thresholded at |t| > 3 and intersected with participant-specific anatomical masks of occipitotemporal brain regions (cuneus, lateral occipital sulcus, pericalcarine gyrus, superior parietal lobule, lingual gyrus, inferior parietal lobule, fusiform gyrus, inferior temporal gyrus, parahippocampal gyrus, middle temporal gyrus), restricted to grey-matter voxels (M = 11,162 voxels on average, SD = 2,083). Classifiers were trained on the volume closest to 4 s after stimulus onset (i.e., the 4th TR, corresponding to a time window of 3.75-5 s post-stimulus), capturing the expected BOLD response peak. Features were detrended per run and standardized (z-scored) separately for each test set.

For subsequent analyses (TDLM, SODA), classifiers trained on all eight runs combined were applied to the sequence trial data, obtaining probabilities for all 13 TRs per trial.

### Probability normalization

For both modalities, decoded classification probability time series were normalized by dividing each class probability by its mean across all time points within a trial. This corrects for baseline offsets between classifiers and ensures that the probability values reflect relative rather than absolute reactivation strength.

### Temporally Delayed Linear Modelling (TDLM)

TDLM quantifies sequential reactivation by testing whether the probability of one stimulus state predicts the probability of the next state in a sequence at a given time lag (Liu et al., 2021). For each sequence trial, the normalized probability time series (5 classes x T time points) were submitted to the TDLM algorithm (Kern et al., 2025). A transition matrix encoding the presented sequence order was constructed. The algorithm computes a “sequenceness” score at each time lag by regressing the probability of the directly following (forward) or preceding (backward) item in the sequence on the probability of the current item, yielding a measure akin to Granger causality for sequential structure. For each trial, 100 control permutations with shuffled transition matrices were computed.

TDLM on MEG. Sequence epochs were extracted starting 200 ms before the first image onset in each sequence trial. For each ISI condition, the analysis window per trial was restricted to the time period during which images were actually shown, to avoid analysing the post-sequence buffer period. The analysis length was computed as ((ISI + 100 ms image duration) x 5 images) / 10 ms sampling interval + max_lag. The maximum tested lag was set to 1.5 times the expected one-step onset-to-onset interval, computed as max_lag = int(((ISI/10) + 10) x 1.5), yielding condition-specific values: 19 time points (190 ms) for the 32 ms ISI, 24 (240 ms) for 64 ms, 34 (340 ms) for 128 ms, and 91 (910 ms) for 512 ms. The factor of 1.5 was chosen to prevent contamination from two-step transitions (e.g., image 1 to image 3) while providing sufficient range at the expected lag. Sequenceness scores were z-scored within each trial across the time lag axis to normalize across participants, as proposed in Wimmer et al. (2020).

TDLM on fMRI. TDLM was applied with a fixed maximum lag of 3 TRs (3.75 s), reflecting the limited temporal resolution of the BOLD signal.

### Slope Order Dynamic Analysis (SODA)

At each measurement time point (TR), the classification probabilities of the five stimuli were regressed onto their serial position in the presented sequence, yielding a regression slope. If stimuli were sequentially reactivated with temporal overlap, slopes are expected to be negative during the early “onset” period (earlier items’ BOLD responses peaking first) and positive during the late “offset” period (later items dominating), as the BOLD responses of earlier items rise and peak before those of later items. Note: following the terminology in the present paper, we refer to these as “onset” and “offset” periods (previously called “forward” and “backward” periods in Wittkuhn & Schuck, 2021) to avoid confusion with forward and backward replay.

SODA on fMRI. We applied SODA to the fMRI sequence trial data (13 TRs per trial). Per convention, SODA slopes were multiplied by-1 to align with the sign conventions of the previous publication. The expected onset and offset periods for each speed condition were derived from the response function parameters fitted by Wittkuhn & Schuck (2021, Table 1): 32 ms ISI: onset TRs 2-4, offset TRs 5-7; 64 ms ISI: onset TRs 2-4, offset TRs 5-7; 128 ms ISI: onset TRs 2-4, offset TRs 5-8; 512 ms ISI: onset TRs 2-5, offset TRs 6-9; 2048 ms ISI: onset TRs 2-7, offset TRs 8-13.

SODA on MEG. Using the same approach as in the fMRI, we regressed the currently shown sequence onto the decoded reactivation probability per time step.

### Group-level statistical testing

TDLM on MEG. For group-level inference, we averaged sequenceness scores across trials per participant and speed condition, and tested for significant sequenceness at each time lag using three statistical approaches:(1) One-sample t-test: A standard one-sample t-test (using scipy, Oliphant, 2007) against zero on the mean sequenceness within a window of +/-10 ms at the expected time, with a significance threshold of p < 0.05. (2) Sign-flip permutation test: A non-parametric permutation t-test (mne.stats.permutation_t_test) with 1024 permutations and t-max correction, testing one-sided (positive sequenceness) at each time lag. A significant result was declared if any time lag within the expected time lag +-10 ms reached p < 0.05. (3) Cluster-based permutation test: A cluster-based permutation test (mne.stats.permutation_cluster_1samp_test) with 512 permutations, which controls for multiple comparisons across time lags by identifying contiguous clusters of significant time points. A significant result required a cluster overlapping with the expected time lag window.

SODA on fMRI. Slopes during the predefined onset and offset periods were tested against zero using analogous statistical approaches: (1) one-sample t-test on the mean slope during each period, (2) sign-flip permutation test with 1024 permutations and t-max correction across all TRs (with offset-period values sign-flipped so both periods are tested as positive effects), and (3) cluster-based permutation test with 512 permutations, where significance requires a cluster overlapping the expected onset or offset TR window. (4) t-test on combined period: We combined onset and offset periods per participant by taking the average per period, flipping the sign of the offset period and comparing the mean of both against zero on a group level using a one-sample t-test.

### Correlation analyses

To assess whether variance in sequenceness was explained by decoder quality or task engagement, we computed correlations between participant-level mean sequenceness across all speed conditions (TDLM) or mean slopes across all speed conditions (SODA) to localizer decoding accuracy at the group average peak and behavioural task performance (proportion of correct position responses) using a linear regression.

### Effect sizes

Effect sizes were quantified using Cohen’s d with 95% confidence intervals (Vallat, 2018, pingouin.compute_effsize). For TDLM, Cohen’s d was computed on the mean sequenceness per participant within the expected time lag window (+/-10 ms) per speed condition. For SODA, Cohen’s d was computed on the mean slope during the onset and offset periods per participant separately.

### Bootstrap power analysis

To estimate the statistical power of TDLM and SODA as a function of sample size and number of trials, we performed a bootstrap resampling analysis. For each combination of sample size (n = 2 to 80) and trial count (n = 2 to 60), we drew bootstrap samples. In each draw, participants were sampled with replacement, and for each sampled participant, the participant’s trials were resampled with replacement. The resampled data were averaged across trials per participant, and the resulting group-level data were tested for significance using each of the three statistical methods described above. Power was defined as the proportion of bootstrap samples yielding a significant result (p < 0.05).

For the t-test variant (1000 draws), significance was assessed by applying a one-sample t-test to the mean sequenceness (TDLM) or mean slope (SODA) within the expected time lag window or TR period or the combined period as defined above. For the sign-flip permutation variant (1000 draws), 1024 sign-flip permutations were computed per bootstrap draw, and significance was declared if any time lag within the expected window (TDLM) or any TR within the expected period (SODA) reached p < 0.05. For the cluster-based variant (512 draws), 512 permutations were computed per draw, and detection required a significant cluster overlapping the expected time lag window or TR period.

### Software

All analyses were implemented in Python. MEG preprocessing and statistical testing was performed using MNE-Python (Gramfort, 2013) and using MEG-BIDS standard (Appelhoff et al., 2019). Machine learning was performed using scikit-learn (Pedregosa et al., 2011). TDLM computations used the TDLM-Python package (Kern et al., 2025). SODA computations used the SODA-Python package (Kern, 2026). fMRI data were organized according to BIDS and accessed via PyBIDS (Yarkoni et al., 2019). Effect sizes were computed using pingouin (Vallat, 2018). Visualizations used matplotlib and seaborn (Hunter, 2007; Waskom, 2021).

## Data and code availability

The code for the analysis and pre-processed probability time series are available at https://github.com/CIMH-Clinical-Psychology/FASTIMAGES-benchmark.

The data for reproduction of this paper are available at https://gin.g-node.org/skjerns/FASTIMAGES-MEG-bids (MEG) https://gin.g-node.org/skjerns/FASTIMAGES-3T-bids (fMRI).

The original fMRI dataset from Wittkuhn and Schuck (2021) is available at https://gin.g-node.org/lnnrtwttkhn/highspeed-bids.

The MEG experiment code can be found at https://github.com/CIMH-Clinical-Psychology/highspeed-task-MEG, the fMRI experiment code can be found at https://github.com/lnnrtwttkhn/highspeed-task.

## Author contributions

SK: Conceptualization, methodology, software, project administration, writing original draft & editing. LW: Conceptualization, methodology, software, reviewing. EB: Investigation, data curation. NS: Conceptualization, supervision, reviewing, resources. GF: Conceptualization, supervision, resources, reviewing.

## Supplement

**Supplement Figure 1.**
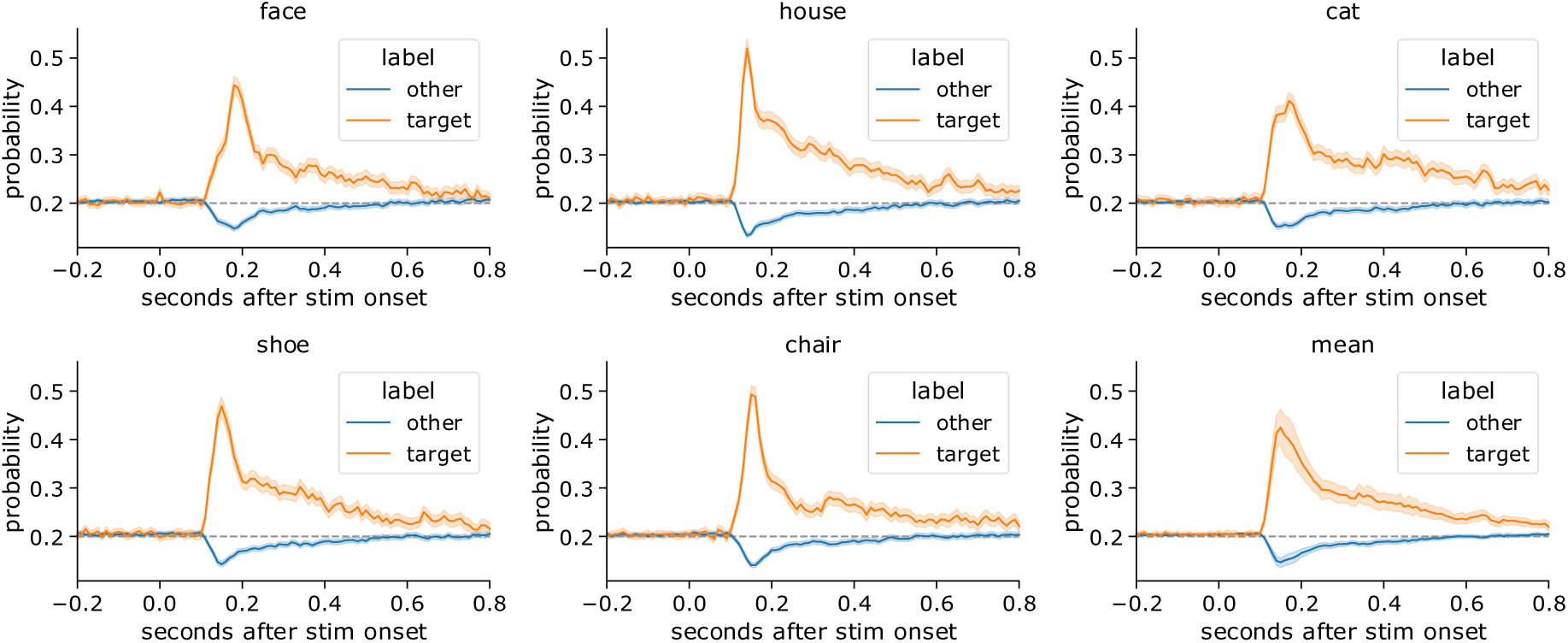
MEG localizer decoding: individual decoder probabilities (one-versus-all logistic regression) during the localizer after image onset, divided by image category.

**Supplement Figure 2.**
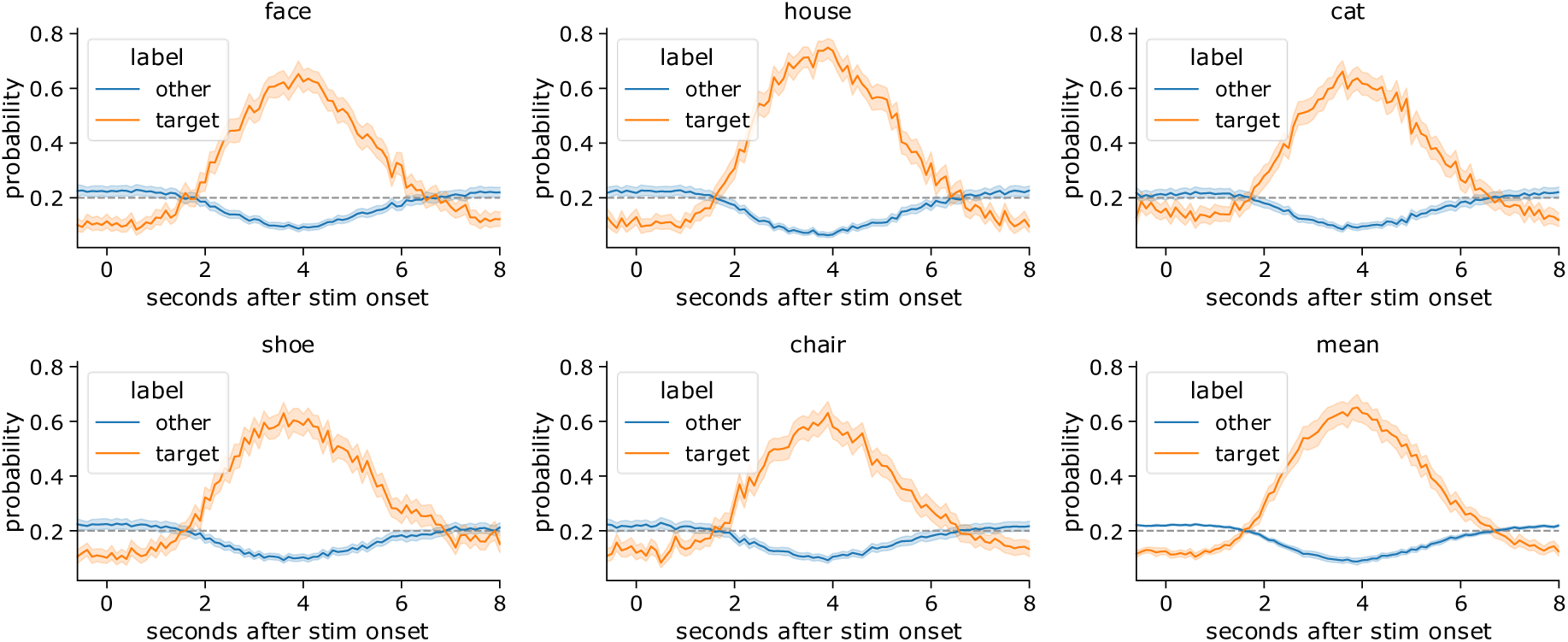
fMRI localizer decoding: individual decoder probabilities (one-versus-all logistic regression) during the localizer after image onset, divided by image category.

**Supplement Figure 3.**
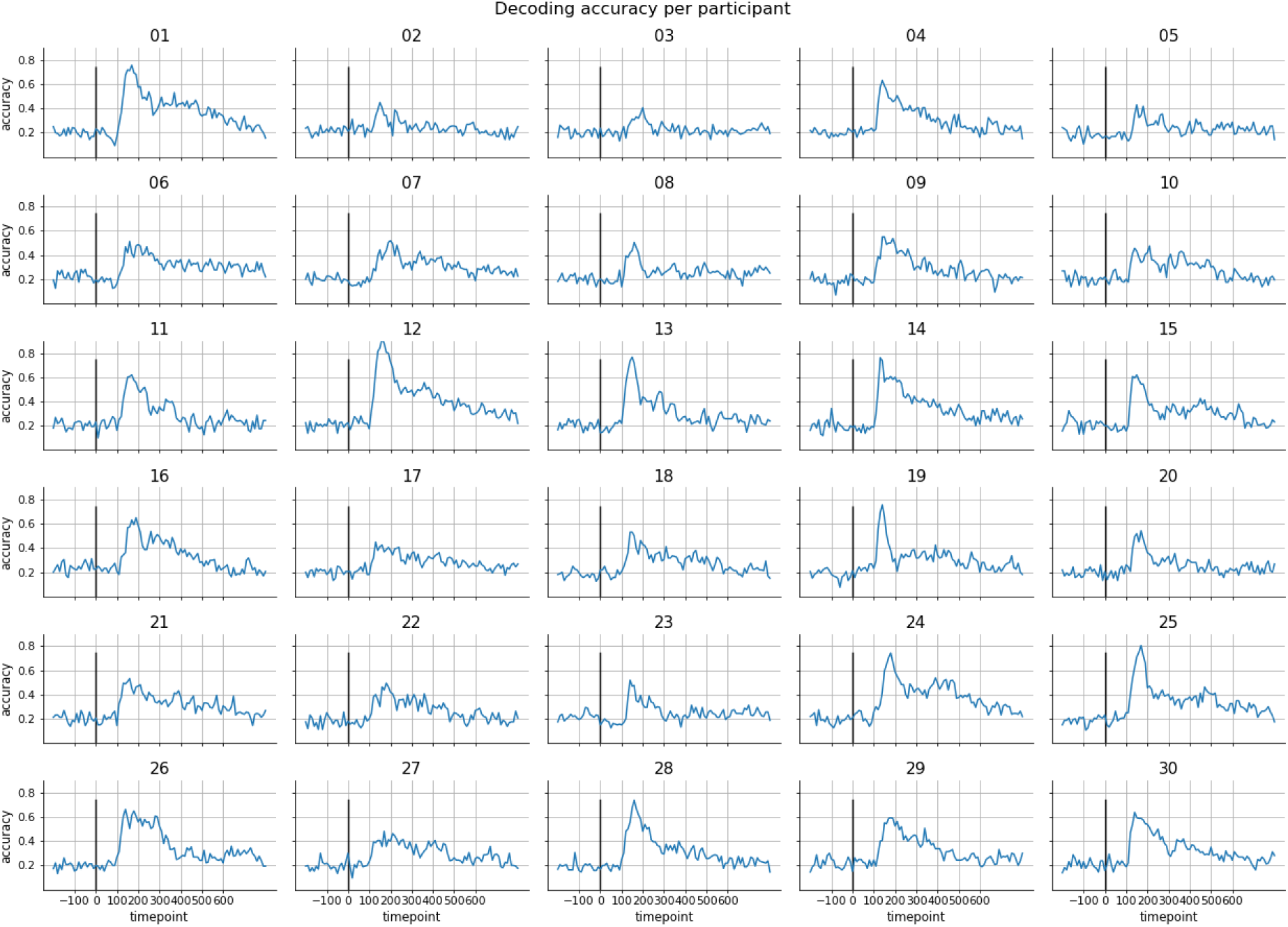
Decoding accuracy per participant in the MEG. Shown is the leave-one-out cross-validated multi-class decoding accuracy. Standard errors are shown in shaded blue bars. A black line indicates the stimulus onset.

**Supplement Figure 4.**
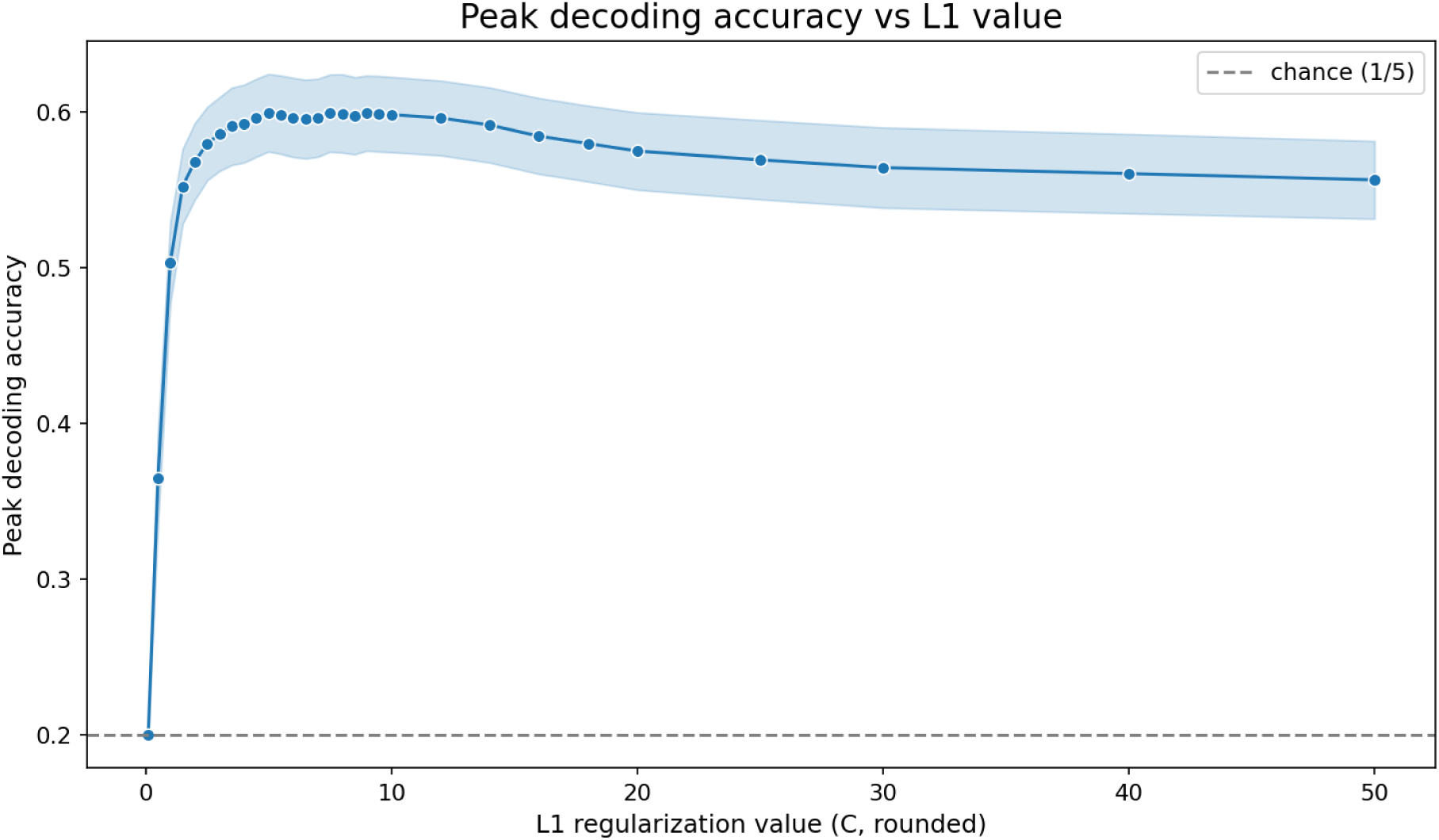
Mean peak decoding accuracy in the MEG for different L1 regularization values C.

**Supplement Figure 5.**
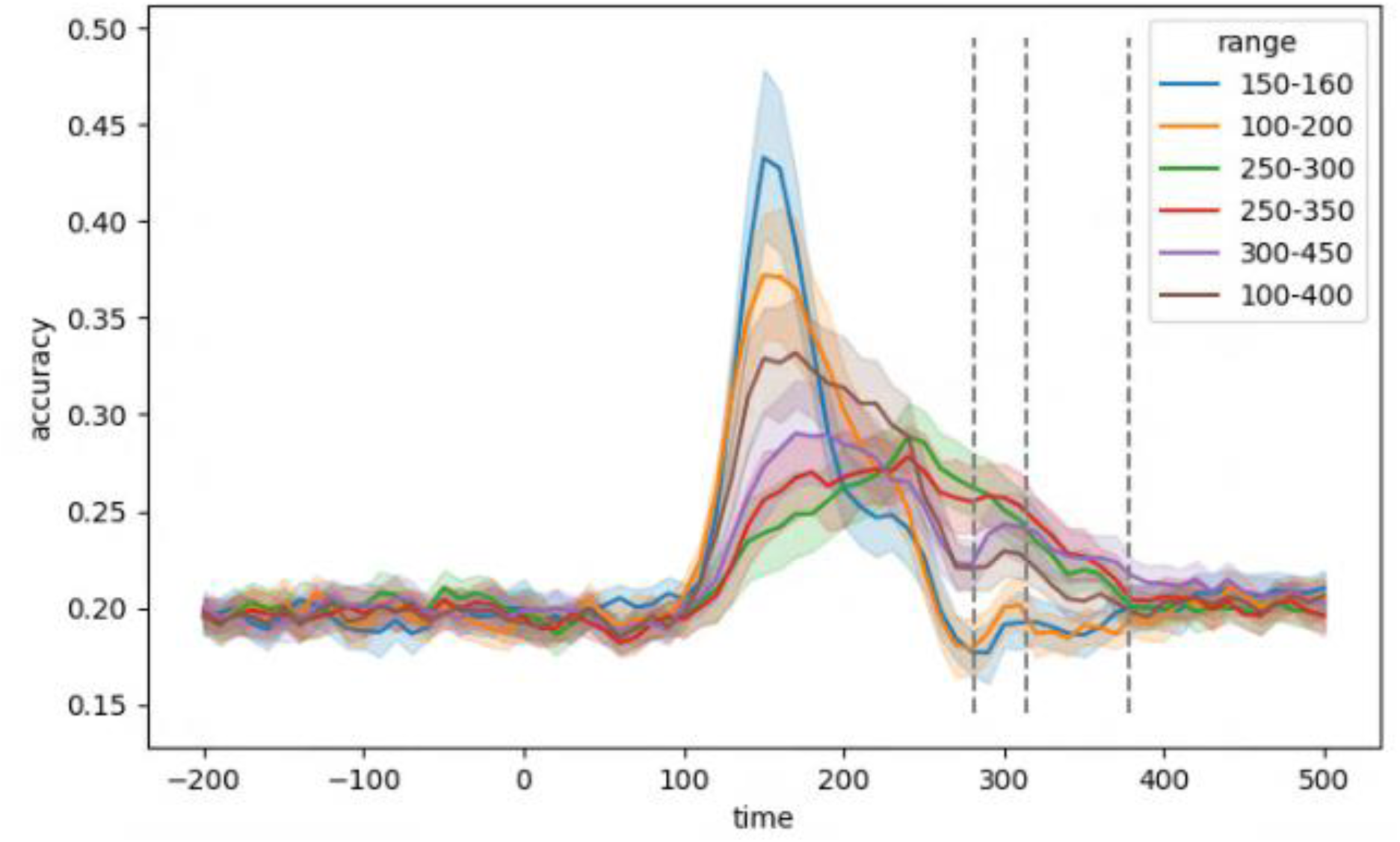
Generalization of classifiers trained on different data segment after stimulus presentation. Usually, classifiers are trained on the peak decoding timepoint. Here, we tried to increasing the breadth of classifier response function by adding data from other timepoints to the training set. While in some cases (e.g., training on 250-350 ms, red line) the response function becomes broader, the decoding accuracy decreases significantly. We assume that the approach of adding several timepoints to the training set creates conflicting information for the classifier. Nevertheless, it is notable, that training on a later timepoint (e.g. after 200 milliseconds) contains information that is not captured by the classifier that is trained on the peak decoding accuracy at 150 milliseconds. It is still unclear which representation is actually being reactivated during replay and whether training on the visual activation peak is capturing this representation well.

**Supplement Figure 6.**
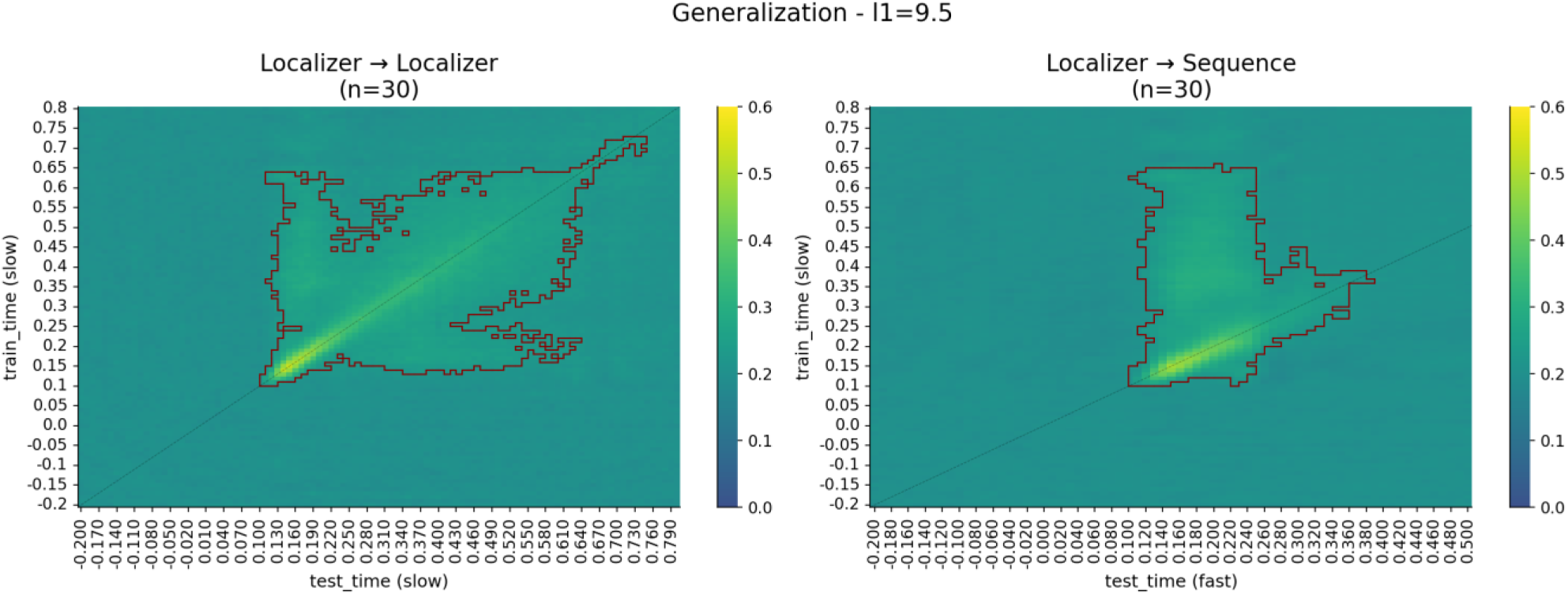
Temporal generalization of the classifier in the MEG. Left: Cross-validated decoding accuracy when training and testing at different time points during the localizer. The decoder trained at the peak decoding accuracy of 150 milliseconds shows a strong generalization towards other time points. Right: A classifier trained on different time points of the localizer, applied to the fast image onsets (all conditions combined). Notably, the image offset seems to stop decodability abruptly, as can be seen by the return to baseline after around 110 milliseconds after processing start, coinciding with the stimulus time of 100 ms. A cluster permutation test was used to determine significance of decoding accuracy above chance, outlined in dark red.

**Supplement Figure 7.**
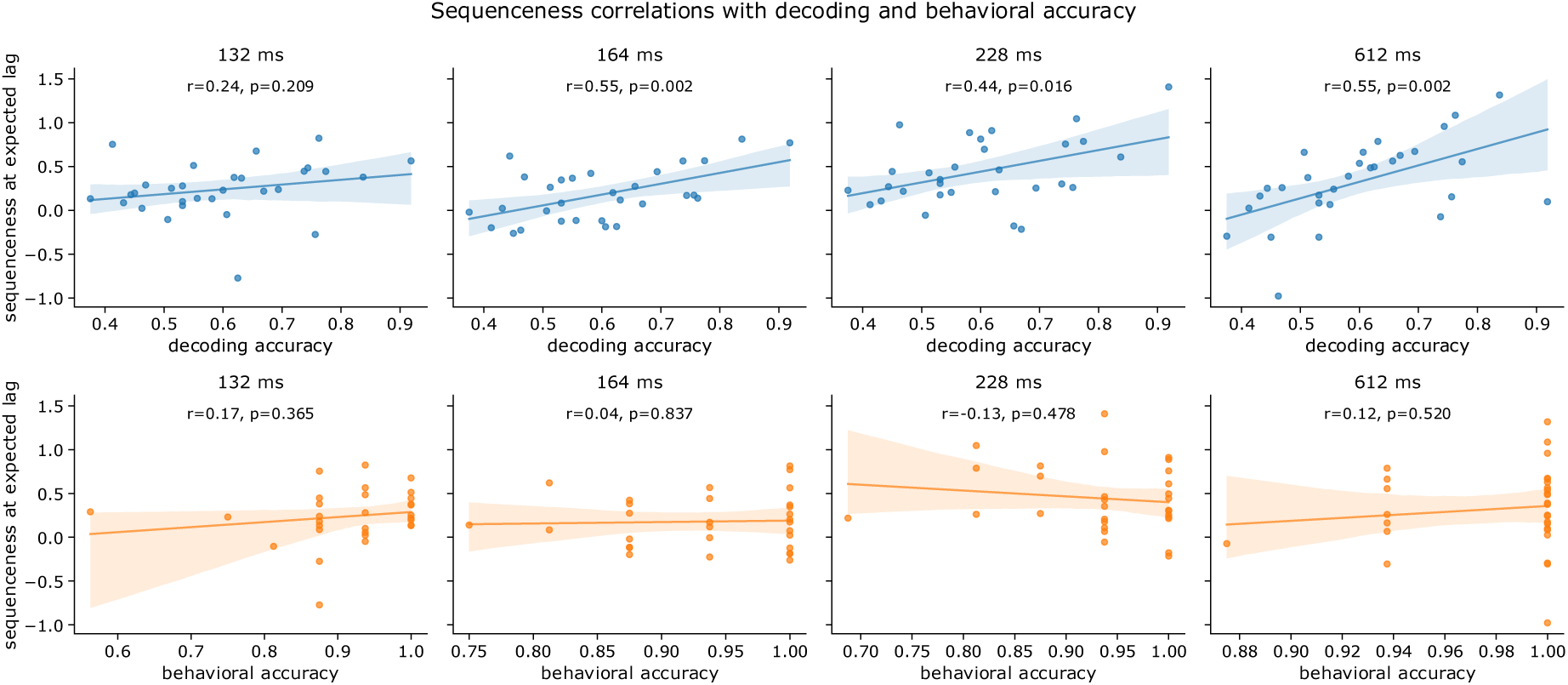
Correlations of mean sequenceness at the expected time lag in the forward direction vs decoding accuracy (upper) and behavioural accuracy (lower) across sequence speeds. Decoding accuracy was significantly related to sequenceness score except in the fastest 132 ms condition. No correlation with behavioural accuracy was present

**Supplement Figure 8.**
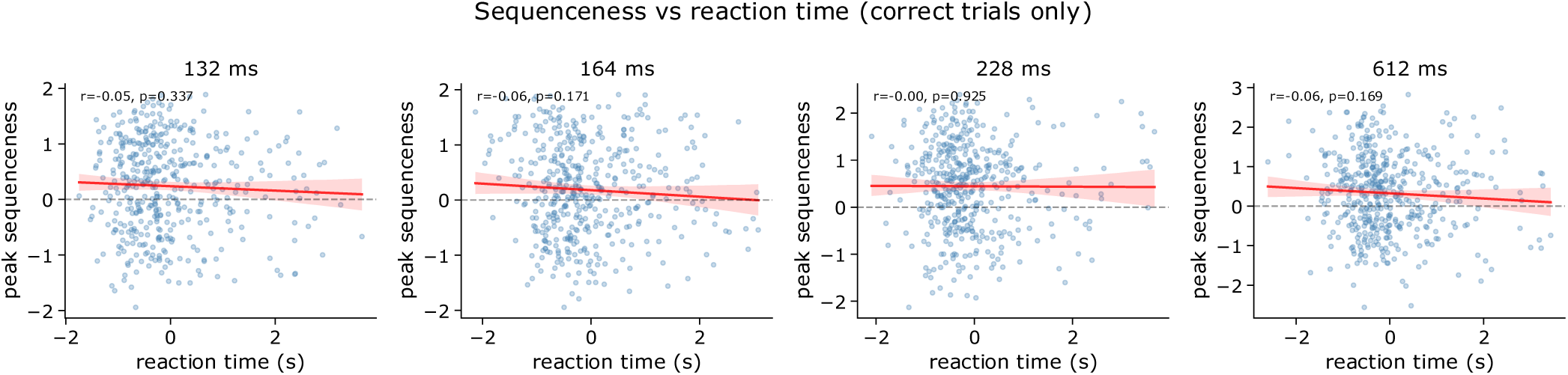
Z-scored reaction times versus peak sequenceness at expected time lag. No interaction between a trials sequenceness and the reaction time was observed.

**Supplement Figure 9.**
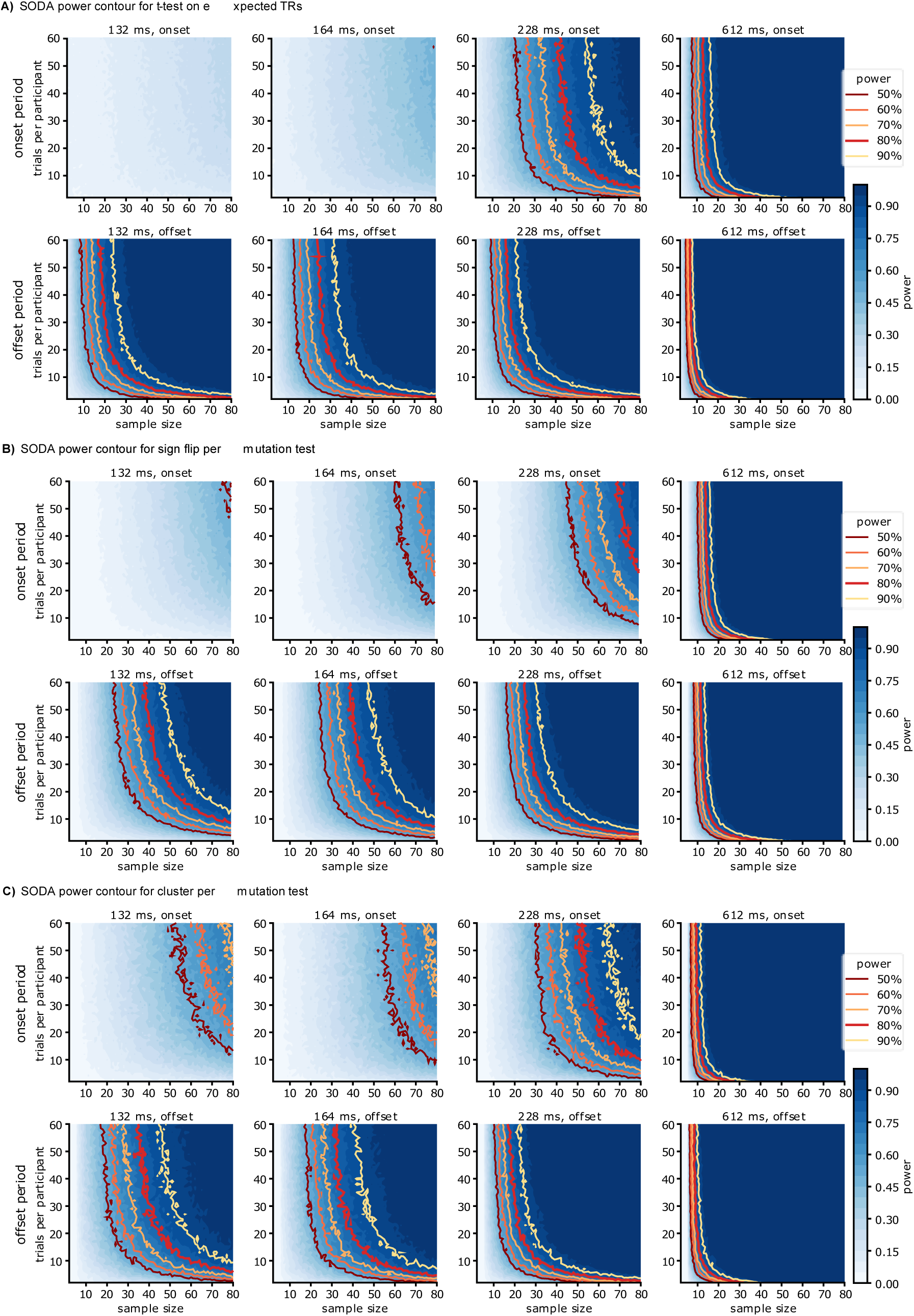
Power contours for SODA using the three different statistical methods. For each combination of sample size and trial size, a bootstrapped sample of that size has been resampled, and per participant the corresponding number of trials has been resampled. We then tested the sample for significance using a **(A)** t-test on the mean slopes during the pre-defined period, or **(B)** a sign-flip permutation test with t-max correction or **(C)** a cluster-permutation test, which was counted as a correct detection if timepoints within the expected time point were significant. We repeated this procedure 1000 times per combination. Resulting power contour plots show what combinations of trial and sample size would have been necessary to detect an effect.

**Supplement Figure 10.**
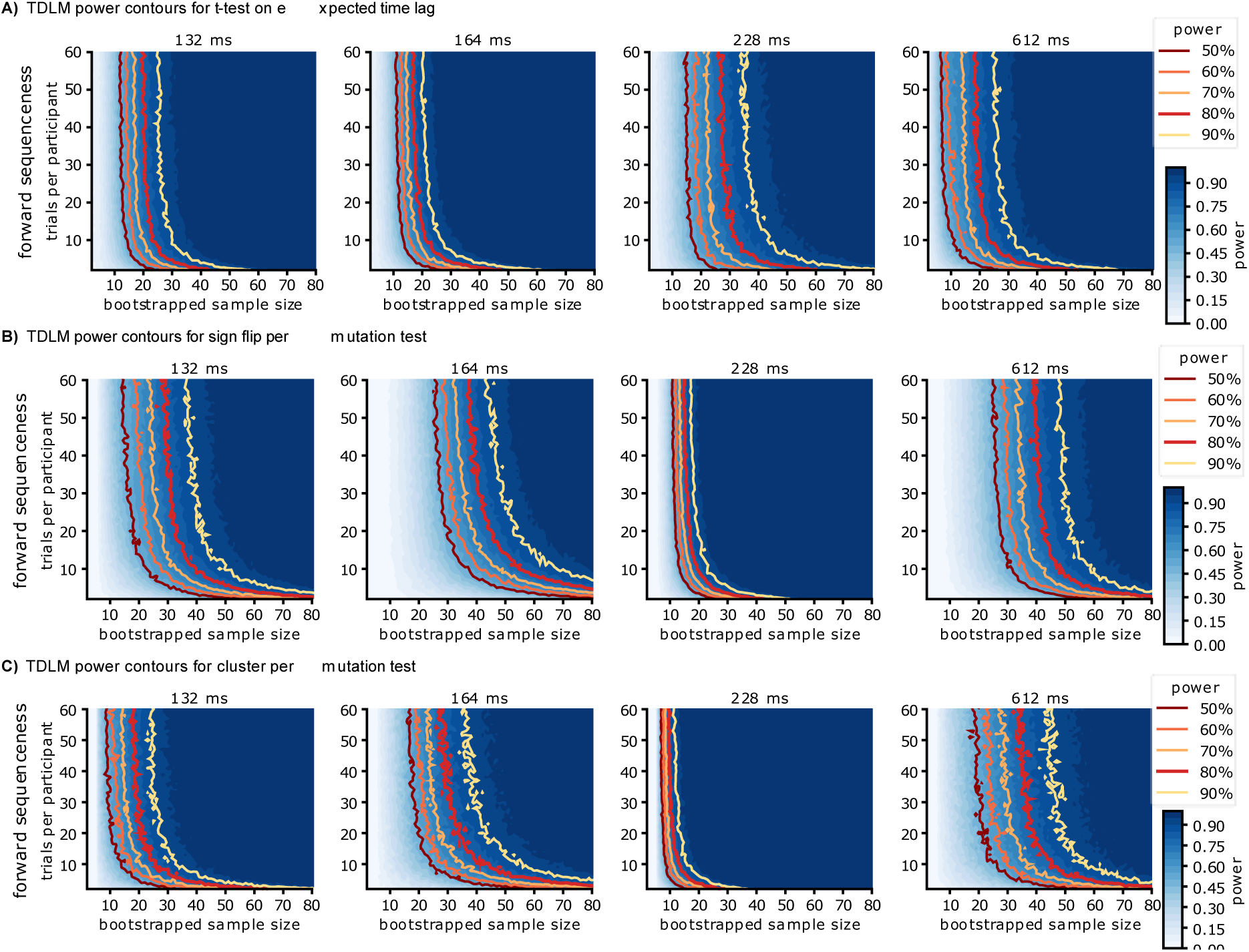
Power contours for TDLM using the three different statistical methods. For each combination of sample size and trial size, a bootstrapped sample of that size has been resampled, and per participant the corresponding number of trials has been resampled. We then tested the sample for significance using a t-test on the mean sequenceness during the expected time lag, or a sign-flip permutation test with t-max correction or a cluster-permutation test, which was counted as a correct detection if timepoints within the expected time lag +-10ms were significant. We repeated this procedure 1000 times per combination. Resulting power contour plots show what combinations of trial and sample size would have been necessary to detect an effect.

**Supplement Figure 11.**
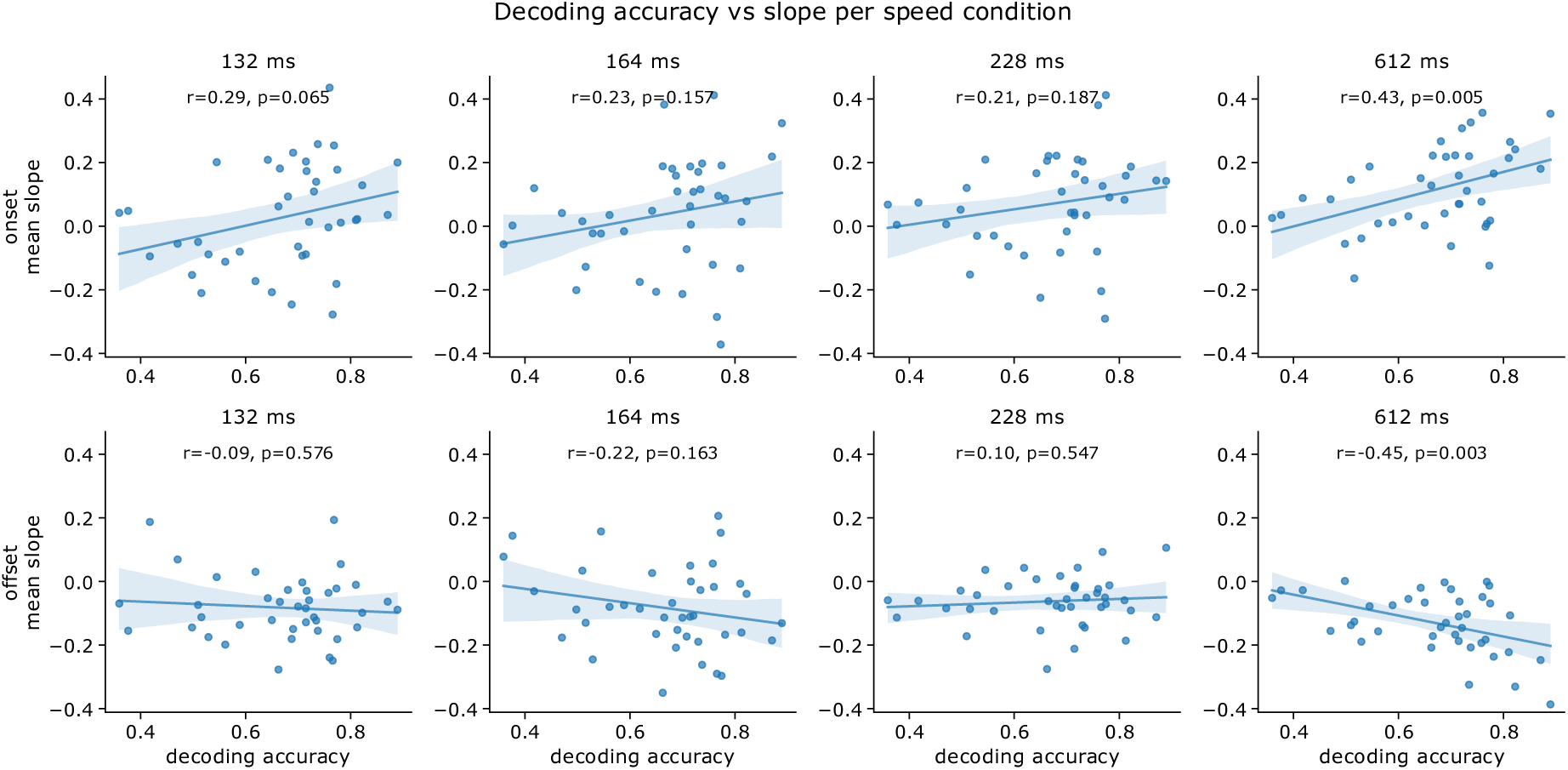
SODA decoding accuracy vs mean slope for onset (upper) and offset (lower) periods.

**Supplement Figure 12.**
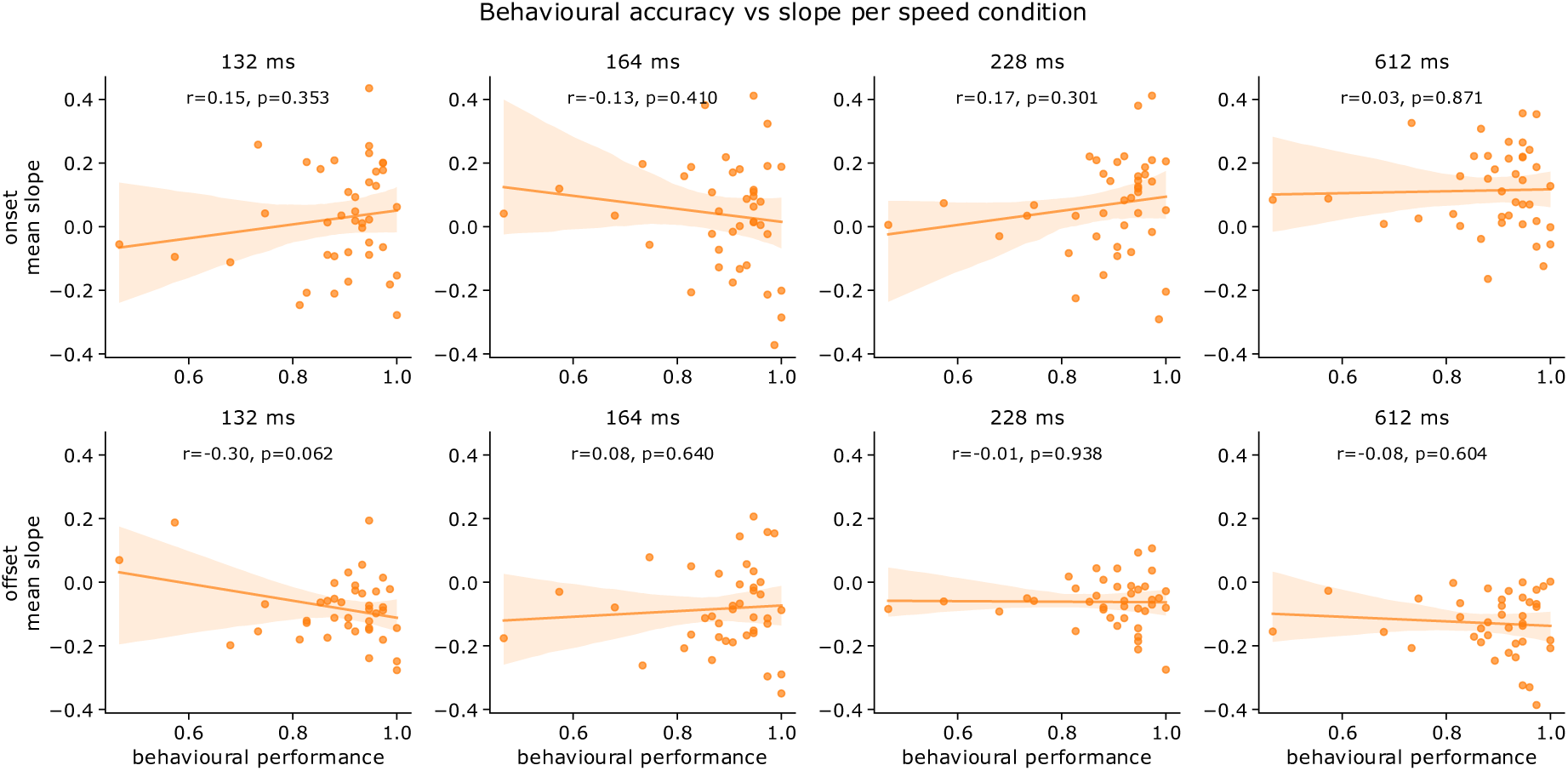
SODA behavioural performance vs mean slope for onset (upper) and offset (lower) periods.

**Supplement Figure 13.**
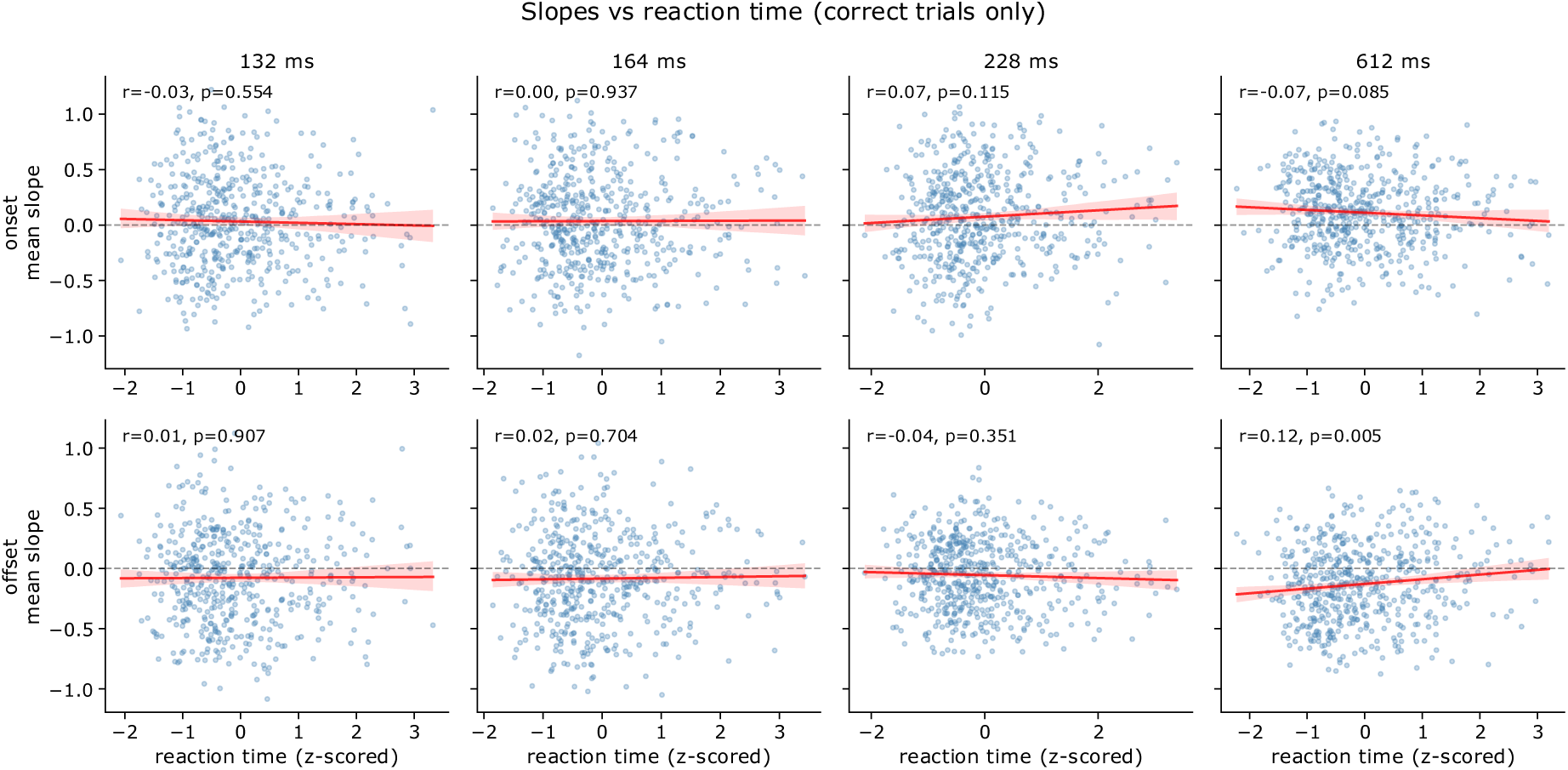
SODA reaction times vs slopes for onset (upper) and offset (lower) period. A significant relationship was present only in the offset of the slowest condition (r=0.12, p=0.005, uncorrected).

